# Targeting Circulating FABP4 Ameliorates Obesity-Associated Hepatic Steatosis

**DOI:** 10.64898/2026.01.24.701451

**Authors:** Xingshan Jiang, Anthony Avellino, Jianyu Yu, Shanshan Liu, Zhaohua Wang, Xiaochun Han, Himani Thakkar, Jonathan Shilyansky, Zhen Xu, Nicholas Schnicker, Bhagirath Chaurasia, Jiaqing Hao, Sonia L. Sugg, Bing Li

## Abstract

Obesity is a major driver of hepatic steatosis, yet the molecular link between excess adiposity and hepatocellular lipid accumulation remains incompletely defined. Here, we identify circulating fatty acid–binding protein 4 (FABP4) as a key mediator of adipocyte–hepatocyte lipid crosstalk in obesity. Analyses of human liver specimens and mouse models reveal aberrant accumulation of FABP4 protein—but not transcript—in hepatocytes during steatosis, indicating an extrinsic source. Genetic deletion of FABP4, specifically in adipocytes, protects against high fat diet-induced hepatic steatosis without altering obesity or systemic lipid levels. Mechanistically, circulating FABP4 directly binds to hepatocytes, facilitating free fatty acid uptake. Furthermore, we developed a high-affinity humanized monoclonal antibody that selectively neutralizes circulating FABP4, blocks hepatocyte binding, suppresses fatty acid uptake, and markedly attenuates hepatic steatosis in multiple obese mouse models. These findings establish circulating FABP4 as a pathogenic lipid chaperone and a promising therapeutic target for obesity-associated hepatic steatosis.

**Graphical Abstract:** 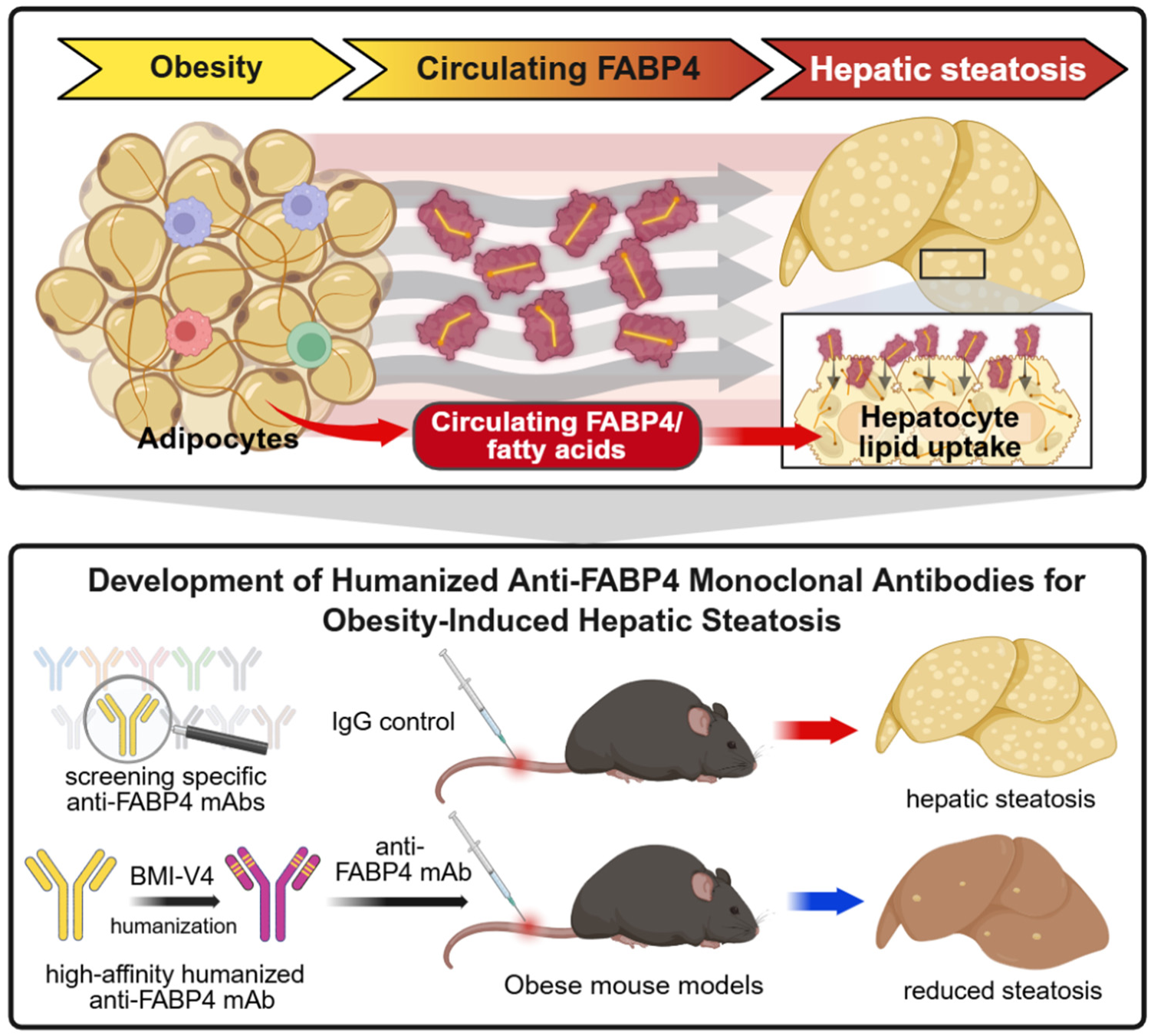

**Highlights:**

1. Hepatocytic accumulation of extrinsic FABP4 links adiposity to liver lipid deposition.
2. Specific deletion of FABP4 in adipocytes prevents hepatic steatosis without affecting systemic lipid levels or obesity.
3. Circulating FABP4 derived from adipocytes directly binds hepatocytes to facilitate free fatty acid transfer.
4. Blocking circulating FABP4 with a high-affinity anti-FABP4 monoclonal antibody inhibits hepatocyte lipid uptake and attenuates steatosis in multiple obese mouse models.

## Introduction

Driven by excess caloric intake and increasingly sedentary lifestyles, obesity has emerged as a global epidemic affecting over 1 billion people worldwide^1^. Obesity is a major risk factor for metabolic syndrome, type 2 diabetes, cardiovascular disease, metabolic dysfunction-associated steatotic liver disease (MASLD), and many types of cancer^2–4^. Among these complications, obesity-associated hepatic steatosis—characterized by excessive triglyceride accumulation in hepatocytes—represents a critical early event in the progression to metabolic dysfunction-associated steatohepatitis (MASH), fibrosis, cirrhosis, and hepatocellular carcinoma^5^.

The pathogenesis of obesity-associated hepatic steatosis is multifactorial. Stable isotope–tracer studies in humans have consistently shown that majority of hepatic lipids in MASLD originate from adipose tissue-derived free fatty acids (FFAs), but not from hepatic *de novo* lipogenesis^6, 7^. In addition, impaired β-oxidation and chronic low-grade inflammation further exacerbate lipid dysregulation and hepatocellular injury^8^. Despite the growing disease burden, current therapeutic options remain limited, consisting primarily of lifestyle modifications and recently approved glucagon-like pepide-1 (GLP-1) receptor agonists (e.g. semaglutide), which provide only modest efficacy and may be accompanied by adverse effects^9, 10^. This unmet clinical need has intensified interest in identifying molecular mediators of adipose–liver crosstalk as potential therapeutic targets.

Among these mediators, adipose fatty acid–binding protein (A-FABP) has emerged as a compelling candidate^11–13^. A-FABP, also known as FABP4 or aP2, is a 15-kDa lipid chaperone predominantly expressed in adipose tissue, including adipocytes, macrophages and endothelial cells^14, 15^. In adipocytes, FABP4 regulates FA trafficking, lipolytic flux, and nuclear receptor signaling. In macrophages, FABP4 promotes inflammatory responses by amplifying NF-κB and JNK signaling and ceramide production, thereby increasing the production of inflammatory cytokines and insulin resistance that accelerate MASLD progression^16–19^. Previous studies were mainly focused on intracellular actions of FABP4 in liver macrophages and endothelial cells^20, 21^, but using small-molecule inhibitors targeting intracellular FABP4 (such as BMS309403) has been hampered in translational studies due to off-target, poor selectivity and severe adverse effects^22–24^. Recently, we and others have demonstrated that obesity induces adipocyte hypertrophy and enhances lipolysis, resulting in increased release of FABP4 into the circulation through unconventional secretion pathways^25–27^. Once in circulation, circulating FABP4 acts as a hormone-like adipokine with systemic metabolic effects that extend beyond adipose tissue^27, 28^. Consistent with this concept, circulating FABP4 levels are strongly associated with obesity, insulin resistance, and MASLD severity in humans^29^, suggesting circulating FABP4 as a previously underappreciated driver and therapeutic target of obesity-associated hepatic steatosis.

In this study, we employed human liver specimens, multiple mouse models (including whole-body and conditional *Fabp4* knockout mice), primary hepatocytes and *Fabp4*-deficient adipocytes to identify adipocyte-derived circulating FABP4 as a principal driver of obesity-associated hepatic steatosis. Given that monoclonal antibody (mAb)-based neutralization offers superior specificity, extended half-life, and the ability to selectively target the pathogenic circulating form of FABP4^30^, we further generated a high-affinity humanized anti-FABP4 mAb and evaluated its therapeutic efficacy in both diet-induced and genetic models of obesity. Collectively, these findings establish precise targeting of circulating FABP4 as an effective strategy to disrupt the adipose–liver axis, reduce hepatic lipid burden, and provide a safe, disease-modifying therapeutic approach for MASLD in the context of obesity.

## Results

### Hepatic steatosis is associated with increased FABP4 protein levels in hepatocytes

When we analyzed publicly available single-cell sequencing datasets from healthy human livers obtained from deceased donors^31, 32^, we noticed that, regardless of donor overweight or obese status (Figure 1A), hepatocytes predominantly express *FABP1*, whereas other *FABP* members, including *FABP4*, were largely absent (Figure 1B, Figure S1A, S1B). These observations indicate that obesity alone does not induce hepatocyte *FABP4* transcription in humans. In contrast, immunohistochemical (IHC) analysis demonstrated that FABP4 protein was undetectable in normal liver tissues but readily detected in hepatocytes from patients with hepatic steatosis and nodular cirrhosis (Figure 1C), with significantly elevated levels compared to normal controls (Figure 1D). These results suggest that FABP4 accumulation in steatotic hepatocytes is unlikely to arise from endogenous transcription and instead may originate from extrinsic sources, contributing to lipid accumulation under pathological conditions.

**Figure 1.**
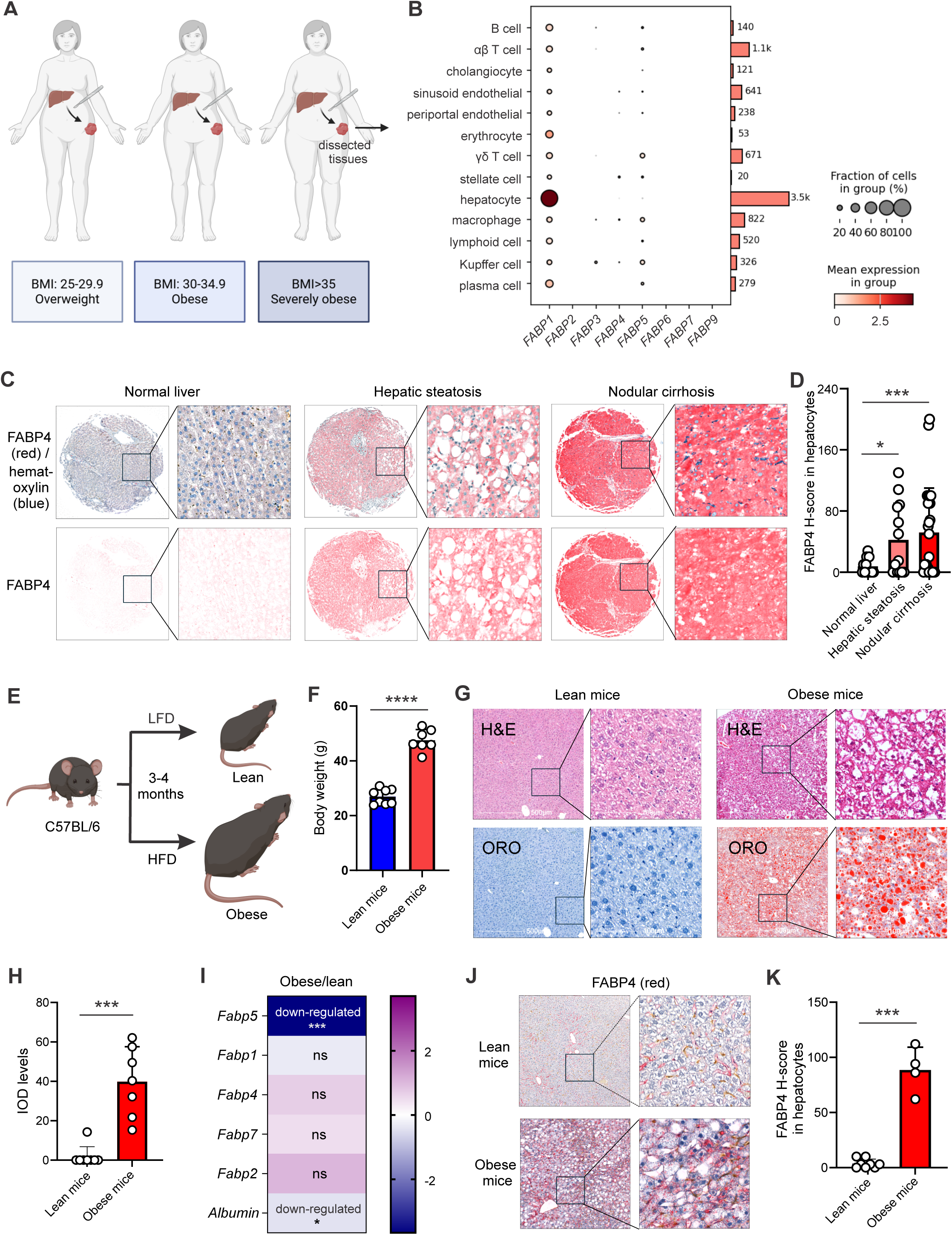
Hepatic steatosis is associated with increased FABP4 protein levels in hepatocytes of both human and mouse livers. **(A)** Healthy liver tissues were obtained from deceased donors with different body mass index (BMI) for single-cell RNA-sequencing analysis. **(B)** Expression profiles of *FABP* family members across single-cell populations in healthy human liver tissues analyzed using BxGenomics. **(C–D)** Representative immunohistochemistry (IHC) staining of FABP4 (red) in human liver tissues from individuals with normal liver, hepatic steatosis, or nodular cirrhosis (C). Quantification of FABP4-positive hepatocytes using H-score is shown in panel **D**. **(E)** Schematic of high-fat diet (HFD)-induced obese mouse models. Low-fat diet (LFD)-fed lean mice served as controls. **(F)** Body weight of obese (HFD-fed) and lean (LFD-fed) mice after 3–4 months of dietary intervention. **(G)** Representative hematoxylin and eosin (H&E) and Oil Red O (ORO) staining of liver tissues from lean and obese mice. **(H)** Quantification of hepatic lipid accumulation based on ORO integrated optical density (IOD). **(I)** Heatmap showing fold changes in hepatic mRNA expression of fatty acid transport-related genes, including *Fabp* family members and *albumin* (*Alb*), in obese mice relative to lean controls. **(L, K)** Representative IHC staining of FABP4 protein accumulation (red) in hepatocytes from lean and obese mice (**J**). Quantification of hepatocytic FABP4 H-score is shown in panel K. Data are shown as mean ± SEM. Statistical significance was determined by Student’s t-test (*, p<0.05, ***p<0.001, ****p<0.0001).

Consistent with the human data, *Fabp4* mRNA was not detected in hepatocytes of normal mouse livers (Figure S1C). In high-fat diet (HFD)-induced obese mouse models (Figure 1E), HFD feeding significantly increased body weight (Figure 1F) and promoted hepatic steatosis, as evidenced by enlarged lipid droplet size (H&E staining) and increased neutral lipid accumulation (Oil Red O staining, ORO) within hepatocytes (Figure 1G, 1H). Importantly, HFD-induced obesity did not significantly increase hepatic *Fabp4* mRNA expression (Figure 1I) or the expression of genes involved in *de novo* fatty acid synthesis and elongation (Figures S1D). Instead, obesity selectively increased FABP4 protein levels in hepatocytes, which strongly correlated with the severity of hepatic steatosis (Figure 1J, 1K). In contrast, FABP5 protein levels in hepatocytes remained low and unchanged in both lean and obese mice (Figure S1E, S1F). Collectively, data from both human and mouse studies demonstrate that hepatocytic accumulation of extrinsic FABP4, rather than transcriptional upregulation, is a defining feature of obesity-associated hepatic steatosis and likely plays a critical role in disease progression.

### FABP4 deficiency attenuates hepatic steatosis in HFD-induced obese mice

To directly assess the role of FABP4 in obesity-associated hepatic steatosis, whole-body *Fabp4* knockout (KO) and wildtype (WT) littermates were fed a HFD to induce obesity and hepatic steatosis (Figure 2A). *Fabp4* deficiency did not reduce HFD-induced body weight gain (Figure 2B) or adipocyte size and morphology compared with WT mice (Figure S2A, S2B), indicating that loss of *Fabp4* does not impair lipid storage under obesogenic conditions. In striking contrast, genetic ablation of *Fabp4* markedly attenuated hepatic steatosis as demonstrated by histological analysis with a substantial reduction of lipid vacuolization in *Fabp4* KO livers. Consistently, ORO staining showed pronounced decreased hepatic neutral lipid accumulation (Figure 2C). Quantitative assessment using ORO staining density further demonstrated that *Fabp4* deficiency significantly reduced obesity-associated lipid deposition in the liver (Figure 2D).

**Figure 2.**
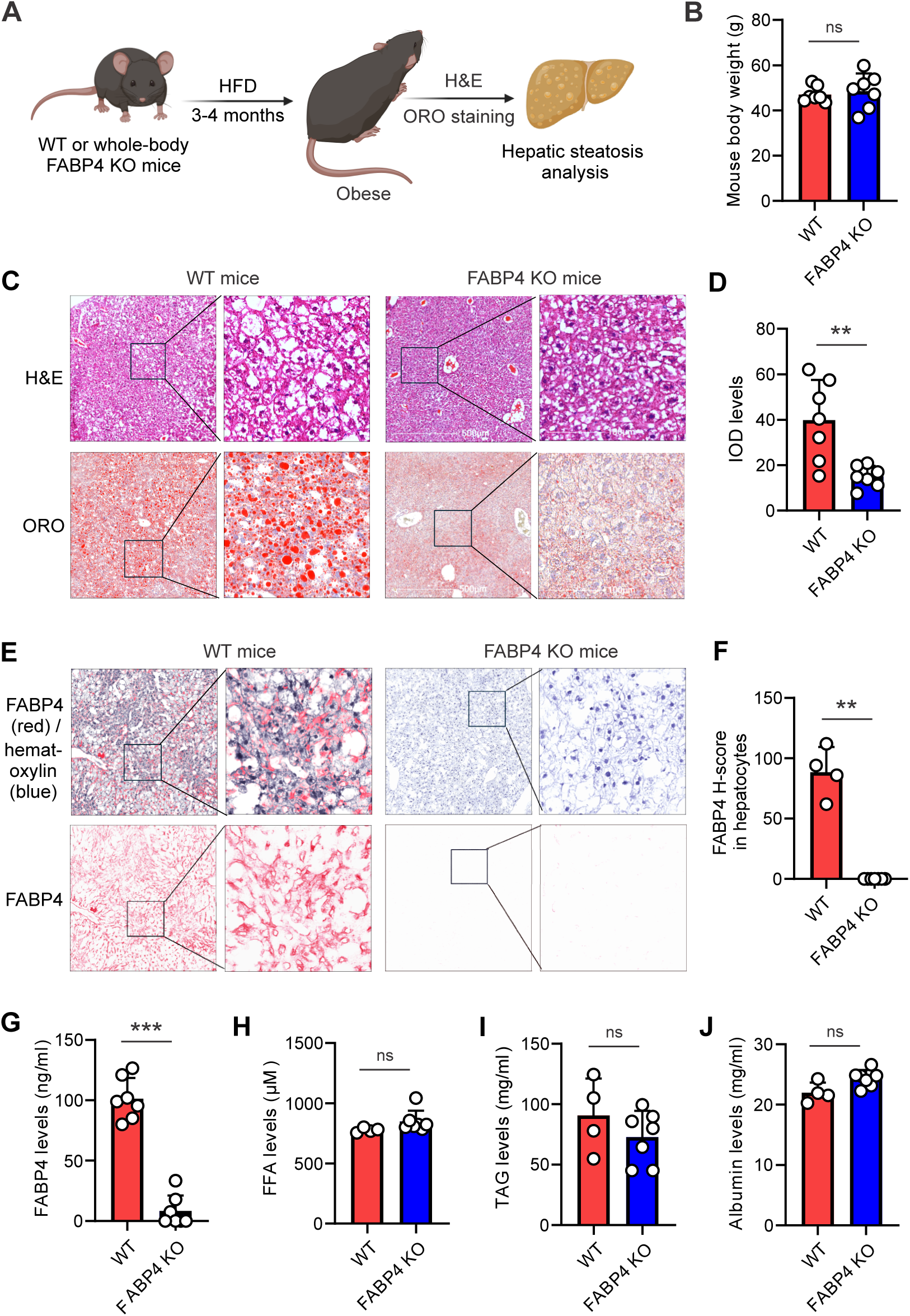
FABP4 deficiency reduces hepatic steatosis in HFD-induced obese mice. **(A)** Schematic illustration of experimental design. Whole-body *Fabp4* knockout (KO) and wild-type (WT) littermates were fed by a high-fat diet (HFD, 60% fat) for 3–4 months to induce obesity, followed by hepatic steatosis analyses. **(B)** Body weight of WT and *Fabp4* KO mice following HFD feeding for 3 months. **(C–D)** Representative hematoxylin and eosin (H&E) and Oil Red O (ORO) staining of liver tissues from obese WT and *Fabp4* KO mice (**C**). Quantification of hepatic lipid accumulation based on ORO integrated optical density (IOD) as shown in panel **D**. **(E–F)** Representative IHC staining of FABP4 protein accumulation (red) in hepatocytes of obese WT and *Fabp4* KO mice (**E**). Quantification of hepatocytic FABP4 H-score is shown in panel **F**. **(G-J)** Analysis of circulating levels of FABP4 (**G**), free fatty acids (FFA) (**H**), triacylglycerol (TAG) (**I**), and albumin (**J**) in serum of obese WT and *Fabp4* KO mice fed with HFD for 3 months. Data are presented as mean ± SEM. Statistical significance was assessed using Student’s t-test (**p<0.01, ***p<0.001, ns, nonsignificant).

Importantly, FABP4 protein was accumulated robustly in hepatocytes of WT mice but was undetectable in *Fabp4* KO livers (Figure 2E, 2F). However, FABP5 protein levels remained low and unchanged in both WT and *Fabp4* KO livers (Figure S2C, S2D), suggesting that the protective phenotype observed in *Fabp4*-defienct mice is unlikely to result from compensatory changes of related *Fabp* family members. Moreover, aside from FABP4 itself, circulating factors associated with lipid availability and transport, including free fatty acids, triglycerides (TAG), and albumin levels, were comparable between WT and *Fabp4* KO mice (Figure 2G-2J). Altogether, these findings identify FABP4 as a nonredundant and essential mediator of obesity-induced lipid accumulation in hepatocytes, demonstrating a direct causal role for FABP4 in the pathogenesis of obesity-induced hepatic steatosis.

### FABP4 specific deletion in adipocytes protects against obesity-associated hepatic steatosis

FABP4 is predominantly expressed in adipocytes, as well as in a subset of macrophages and endothelial cells^33^. To define the cell type-specific contribution of FABP4 to obesity-associated hepatic steatosis, we generated *Fabp4* floxed mice (*Fabp4*^f/f^) by flanking exons 2 and 3 of the *Fabp4* gene with LoxP sites (Figure S3A). *Fabp4*^f/f^ mice were subsequently crossed with Adipoq-Cre, CSF1R-Cre, or Tek-Cre transgenic lines to generate adipocyte-, macrophage-, or endothelial cell-specific *Fabp4* knockout mice, respectively (Figure 3A).

**Figure 3.**
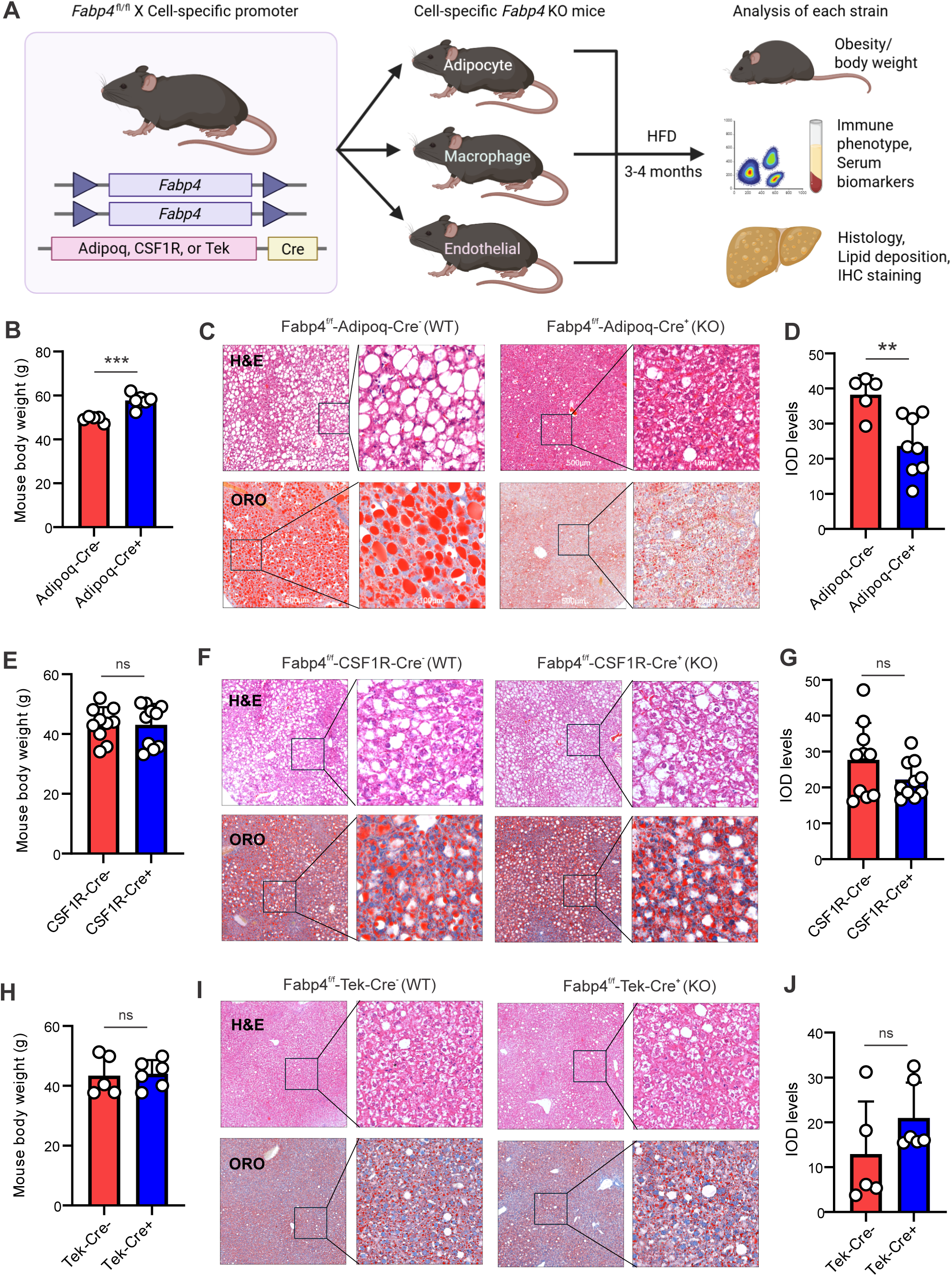
Adipocyte-specific deletion of FABP4 attenuates HFD-induced hepatic steatosis. **(A)** Schematic of the experimental design. *Fabp4*^f/f^ mice were crossed with Adipoq-Cre, CSF1R-Cre, or Tek-Cre drivers to generate adipocyte-, macrophage-, or endothelial cell–specific FABP4 knockout mice. Mice were fed a high-fat diet (HFD) for 3–4 months, followed by assessment of metabolic parameters and hepatic steatosis. **(B)** Body weight of *Fabp4*^f/f^ Adipoq-Cre- (WT) mice and Cre+ (KO) controls after HFD feeding. **(C)** Representative hematoxylin and eosin (H&E) and Oil Red O (ORO) staining of liver tissues from *Fabp4*^f/f^ Adipoq-Cre- (WT) mice and Cre+ (KO) mice following HFD feeding. **(D)** Quantification of hepatic lipid accumulation based on ORO integrated optical density (IOD) as shown in panel **C**. **(E)** Body weight of *Fabp4*^f/f^ CSF1R-Cre- (WT) mice and Cre+ (KO) mice following HFD feeding for 3 months. **(F)** Representative H&E and ORO staining of liver tissues from *Fabp4*^f/f^ CSF1R-Cre- (WT) mice and Cre+ (KO) mice following HFD feeding. **(G)** Quantification of hepatic lipid accumulation based on ORO integrated optical density (IOD) as shown in panel F. **(H)** Body weight of *Fabp4*^f/f^ TEK-Cre- (WT) mice and Cre+ (KO) mice after HFD feeding. **(I)** Representative H&E- and ORO-stained liver sections *Fabp4*^f/f^ TEK-Cre- (WT) mice and Cre+ (KO) mice following HFD feeding. **(J)** Quantification of hepatic lipid accumulation based on ORO integrated optical density (IOD) as shown in panel I. Data are presented as mean ± SEM. Statistical significance was determined by Student’s *t*-test (**,*p*<0.01; ***,*p*<0.001; ns, nonsignificant).

To validate the efficiency and specificity of *Fabp4* deletion, FABP4 expression was assessed in adipocytes, F4/80^+^ macrophages, and CD31^+^ endothelial cells. In *Fabp4*^f/f^ Adipoq-Cre mice, IHC analysis demonstrated selective deletion of FABP4, but not FABP5, in adipose tissue (Figure S3B-S3C). Intracellular flow cytometry further confirmed efficient FABP4 depletion in adipocytes with no detectable loss in CD45^+^ immune cells (Figure S3D-S3E). In addition, *Fabp4*^f/f^ CSF1R-Cre and *Fabp4*^f/f^ Tek-Cre mice showed efficient and specific deletion in macrophages (Figure S3F, S3G) and endothelial cells, respectively (Figures S3H, S3I), confirming robust cell type-specific *Fabp4* knockout across these models.

We next assessed the functional contribution of cell type-specific FABP4 deletion to HFD-induced obesity and hepatic steatosis (Figure 3A). After 3 months of HFD feeding, *Fabp4*^f/f^ Adipoq-Cre^+^ mice exhibited greater body weight than *Fabp4*^f/f^ Adipoq-Cre^-^ littermates (Figure 3B), consistent with impaired adipocyte lipolysis and enhanced adiposity following adipocyte-specific FABP4 deletion. Despite increased adiposity, histological analysis by H&E staining revealed markedly fewer and smaller lipid vacuoles and preservation of hepatic architecture in Adipoq-Cre^+^ livers compared with Adipoq-Cre^-^ controls (Figure 3C). Consistently, ORO staining showed a substantial reduction in neutral lipid accumulation in Adipoq-Cre⁺ livers, as confirmed by quantitative analysis (Figure 3D). In contrast, FABP4 deletion in macrophages (Figure 3E-3G) or endothelial cells (Figure 3H-3J) neither altered HFD-induced obesity, nor affected obesity-associated hepatic steatosis, suggesting that FABP4 in these cell types is dispensable for obesity-induced steatosis development. Altogether, these findings demonstrate that adipocyte-derived FABP4, rather than macrophage- or endothelial-derived FABP4, is a key driver of obesity-associated hepatic lipid accumulation, highlighting a previously underappreciated adipocyte–hepatocyte crosstalk mechanism in hepatic steatosis.

### Circulating FABP4 facilitates free FA uptake by hepatocytes

To elucidate the mechanism by which adipocyte-derived FABP4 promotes hepatic steatosis, we observed that FABP4 was able to be secreted out into the circulation in WT mice but was undetectable in the circulation of whole-body *Fabp4* KO mice (Figure 4A). Circulating FABP4 levels were significantly reduced in adipocyte-specific *Fabp4* KO mice but remained unchanged in macrophage- or endothelia cell-specific *Fabp4* KO mice (Figure 4B-4D), indicating that adipocytes are the primary source of circulating FABP4. Furthermore, Adipoq-Cre^+^ mice exhibited reduced FABP4 accumulation in hepatocytes compared to Adipoq-Cre^-^ mice (Figure S4A, S4B), suggesting an adipocyte origin of liver FABP4. Based on these observations, we hypothesized that circulating FABP4 derived from adipocytes facilitates lipid transport and hepatocyte uptake. To test this, hepatocyte cell lines (Huh7 and Hepa1-6) were incubated with fluorescently labeled free fatty acids (FAs), including oleic acid (TopFluor^TM^-OA) and palmitic acid (Bodipy^TM^ FL C-16, PA), in the presence or absence of soluble FABP4. In the absence of FABP4, uptake of free OA by both Huh7 and Hepa1-6 cells was minimal. Addition of FABP4 markedly enhanced OA uptake in both cell lines (Figure 4E-4H). Moreover, FABP4 promoted OA uptake by hepatocyte in a dose-dependent manner (Figure S4C-S4D). Consistent with this effect, FABP4 also significantly increased PA uptake in both hepatocyte models (Figure S4E-S4H). These data demonstrate that FABP4 is sufficient to facilitate hepatocellular uptake of both saturated and unsaturated FAs.

**Figure 4.**
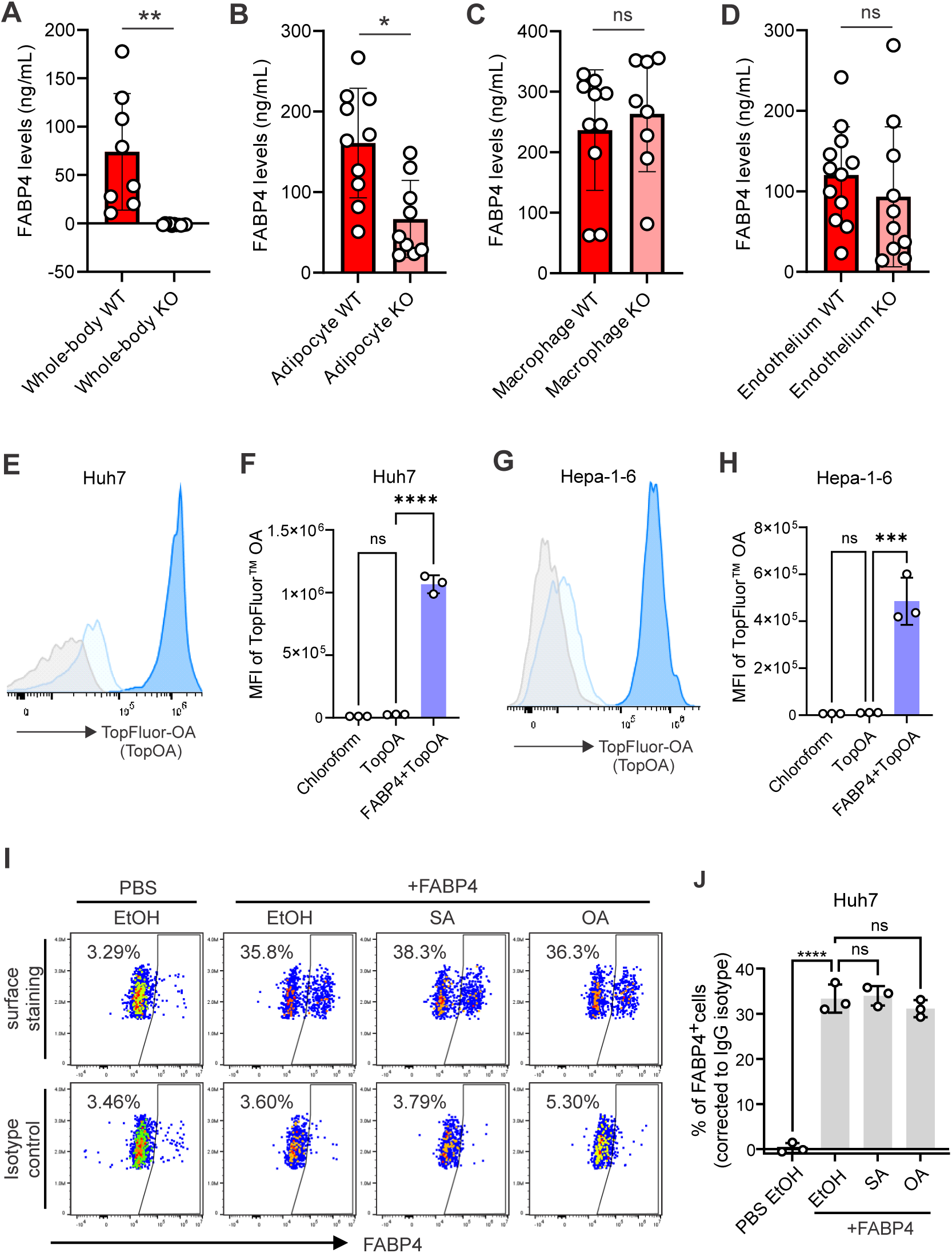
Circulating FABP4 binds to the cell surface and mediates free fatty acids transfer to HCCs. **(A–D)** Serum concentrations of FABP4 in HFD-fed obese mice with whole-body *Fabp4* knockout (*Fabp4* KO)(A), adipocyte-specific *Fabp4* KO (Adipoq-Cre *Fabp4*^f/f^) (B), macrophage-specific *Fabp4* KO (CSF1R-Cre *Fabp4*^f/f^) (C), or endothelial-specific *Fabp4* KO (Tek-Cre *Fabp4*^f/f^) (D) as measured by ELISA. **(E–H)** Representative flow cytometric histogram (E, G) and Quantification of TopFluor^TM^ OA fluorescence intensity (F, H) in Huh7(E - F) and Hepa1-6 (G – H) after 15 min incubation with either free form or FABP4-bound form (1:1) of TopFluor^TM^ OA (1 μmol/L). Chloroform was used as a vehicle control. **(I–J)** Representative flow cytometric gating (I) and quantification (J) of FABP4 on the surface of Huh7 after 15 min incubation of FABP4-bound free fatty acid (1:1), including either SA or OA at 1 μmol/L concentration. EtOH were used as vehicle control. Data are presented as mean ± SEM. Statistical significance determined by unpaired two-tailed t-test or one-way ANOVA. ***, p < 0.001; ****, p < 0.0001; ns, nonsignificant.

While assessing Huh7 hepatocytes, we noticed that, consistent with our *in vivo* data, these hepatocytes did not endogenously express FABP4 (Figure 4I, 4J). However, when exposed to soluble FABP4, either FA loaded or unloaded, FABP4 directly bound to the surface of a subset of Huh7 cells, thereby enabling efficient FA transfer. Similar surface binding of FABP4 was also observed in Hepa1-6 cells (Figure S4I-S4J), suggesting a common mechanism across different hepatocytes. Taken together, these results identify that circulating FABP4 directly binds to hepatocytes and functions as a carrier to facilitate free FA uptake, providing a mechanistic basis for its causal role in obesity-associated hepatic steatosis.

### Circulating FABP4 mediates adipocyte-hepatocyte lipid crosstalk

To establish the causal role for circulating FABP4 in adipocyte-hepatocyte lipid crosstalk, we generated stable *Fabp4* KO 3T3 preadipocyte cell lines using CRISPR/Cas9 technology (Figure S5A). Upon adipogenic differentiation, FABP4 was highly expressed in mature WT 3T3 adipocytes but was completely absent in FABP4 KO 3T3 cells (Figure 5A, 5B).

**Figure 5.**
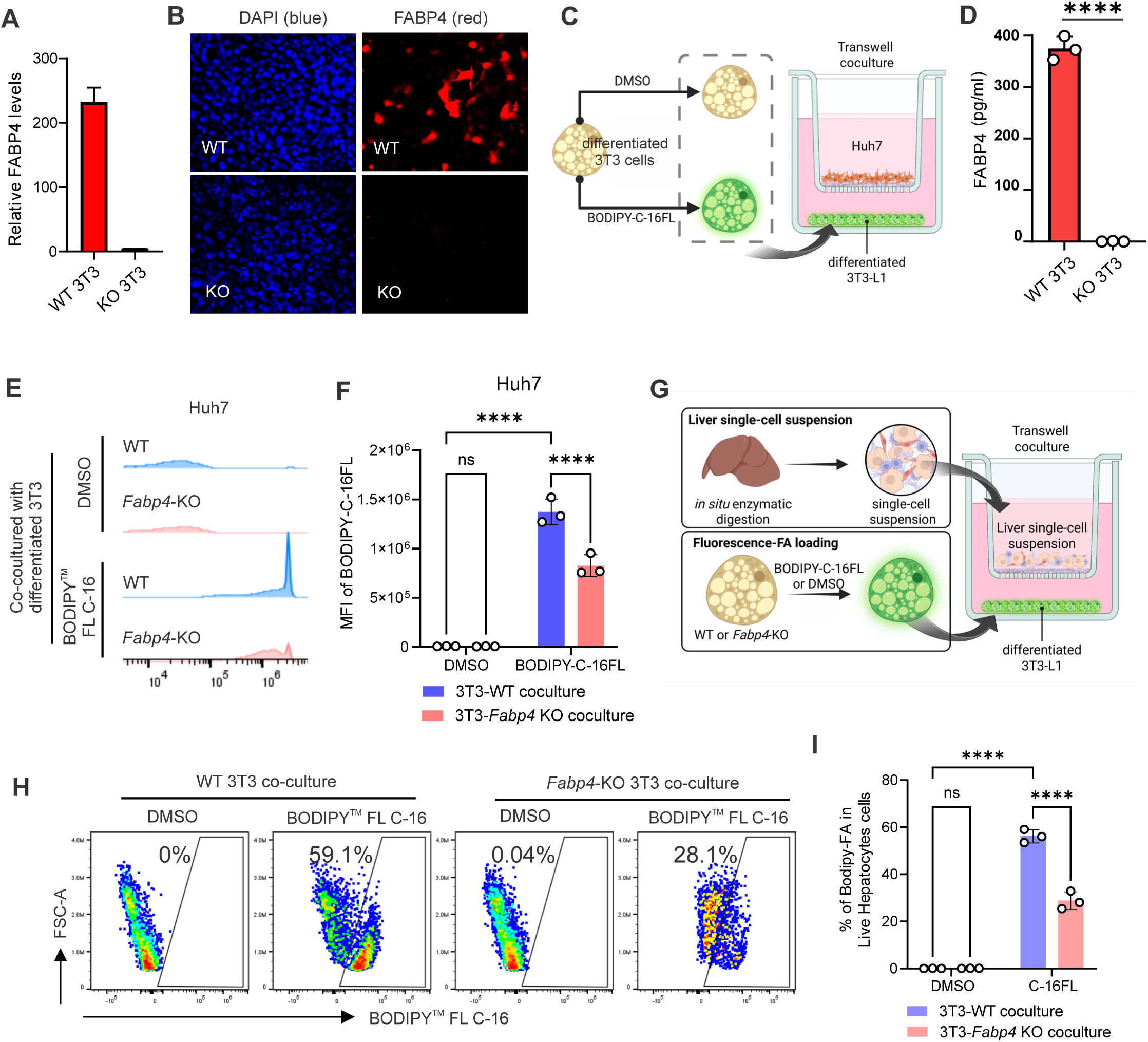
Adipocyte-derived FABP4 mediates lipid transfer to hepatocytes. **(A–B)** Validation of *Fabp4*-KO in differentiated 3T3 cells via qPCR (A) and immunofluorescence staining (B) showing ablated level of *Fabp4* expression in differentiated *Fabp4*-KO 3T3 cells. **(C)** Schematic of *in vitro* transwell fluorescent-labelled FA transfer of Huh7 in the insert, cocultured with either DMSO or BODIPY^TM^ FL C-16-loaded differentiated 3T3 cells in the low chamber. **(D)** ELISA measurement of FABP4 in the supernatant collected after 16 h culture of differentiated WT or *Fabp4*-KO 3T3 cells in RPMI. **(E–F)** Representative flow cytometric histogram (E) and Quantification of BODIPY^TM^ FL C-16 fluorescence intensity (F) in Huh7 after 24 h of coculture with either WT or *Fabp4*-KO differentiated 3T3, which were initially loaded with BODIPY^TM^ FL C-16 (1 μmol/L). DMSO was used as a vehicle control in both WT and *Fabp4*-KO differentiated 3T3. **(G)** Schematic of transwell fluorescent-labelled FA transfer of liver primary cells in the insert, cocultured with either DMSO or BODIPY^TM^ FL C-16-loaded differentiated 3T3 cells in the lower chamber. **(H–I)** Representative flow cytometric gating (H) and Quantification of BODIPY^TM^ FL C-16 fluorescence intensity (I) in primary hepatocytes after 24 h of coculture with either WT or *Fabp4*-KO differentiated 3T3, which were initially loaded with BODIPY^TM^ FL C-16 (1 μmol/L). DMSO was used as a vehicle control in both WT and *Fabp4*-KO differentiated 3T3. Data are presented as mean ± SEM. Statistical analysis was performed using unpaired t-tests or two-way ANOVA. ****, p < 0.0001; ns, nonsignificant.

To model adipocyte-hepatocyte lipid communication *in vitro*, fluorescent PA-loaded *Fabp4* WT or KO adipocytes were cocultured with Huh7 hepatocytes using a transwell system (Figure 5C). FABP4 was readily secreted into the culture supernatant by WT adipocytes (Figure 5D) and efficiently mediated the transfer of PA to Huh7 cells (Figure 5E). In contrast, genetic ablation of *Fabp4* in 3T3 adipocytes significantly impaired PA transfer to Huh7 cells (Figure 5F). Similarly, *Fabp4* deficiency in adipocytes also reduced PA transfer to Hepa1-6 cells (Figure S5B, S5C), demonstrating that adipocyte-derived FABP4 is required for efficient FA transfer to hepatocytes.

To extend these findings to a physiologically relevant system, primary cells were isolated from mouse livers using an *in situ* enzymatic digestion protocol (Figure S5D). Primary hepatocytes accounted for over 90% of liver cell populations (Figure S5E) and were cocultured with fluorescent PA-loaded *Fabp4* WT or KO adipocytes in a transwell system (Figure 5G). Consistent with the hepatocyte cell line data, *Fabp4* deficiency in adipocytes markedly reduced PA transfer to primary hepatocytes compared with WT controls (Figure 5H, 5I). Notably, adipocyte-derived FABP4 preferentially bound to primary hepatocytes relative to other hepatic cell types (Figure S5F), and primary hepatocytes exhibited the highest PA uptake among liver cells (Figure S5G).

Altogether, these results demonstrate that adipocyte-derived circulating FABP4 facilitates FA transfer and uptake by hepatocytes in both cell line-based and primary cell systems, establishing an underappreciated causal role for circulating FABP4 in adipocyte-hepatocyte lipid crosstalk underlying hepatic steatosis.

### Generation of a high-affinity humanized anti-FABP4 antibody to block circulating FABP4 activity

Given our findings that circulating FABP4 mediates lipid crosstalk between adipocytes and hepatocytes, we next investigated whether circulating FABP4 activity could be therapeutically blocked. Through a comprehensive screen for neutralizing anti-FABP4 monoclonal antibodies (mAb) across different animal models^30^, we identified a clone, termed BMI-V4, that selectively bound FABP4 but not to other FABP members, including FABP5 (Figure 6A). Mass photometry analysis confirmed specific, dose-dependent binding of BMI-V4 to FABP4 (Figure 6B, S6A-S6C). Notably, compared with previously reported anti-FABP4 antibodies with relatively low affinity (e.g., CA33, Kd of the μM range^34^), BMI-V4 exhibited exceptionally high binding affinity, with a dissociation constant in the picomolar range (Figure 6C). To enhance translational potential, we generated a humanized version of the V4 antibody by CDR-grafting onto the most homologous human germline frameworks, as previously described^30^. Recombinant humanized V4 antibody was purified in CHO cells for functional analyses (Figure S6D).

**Figure 6.**
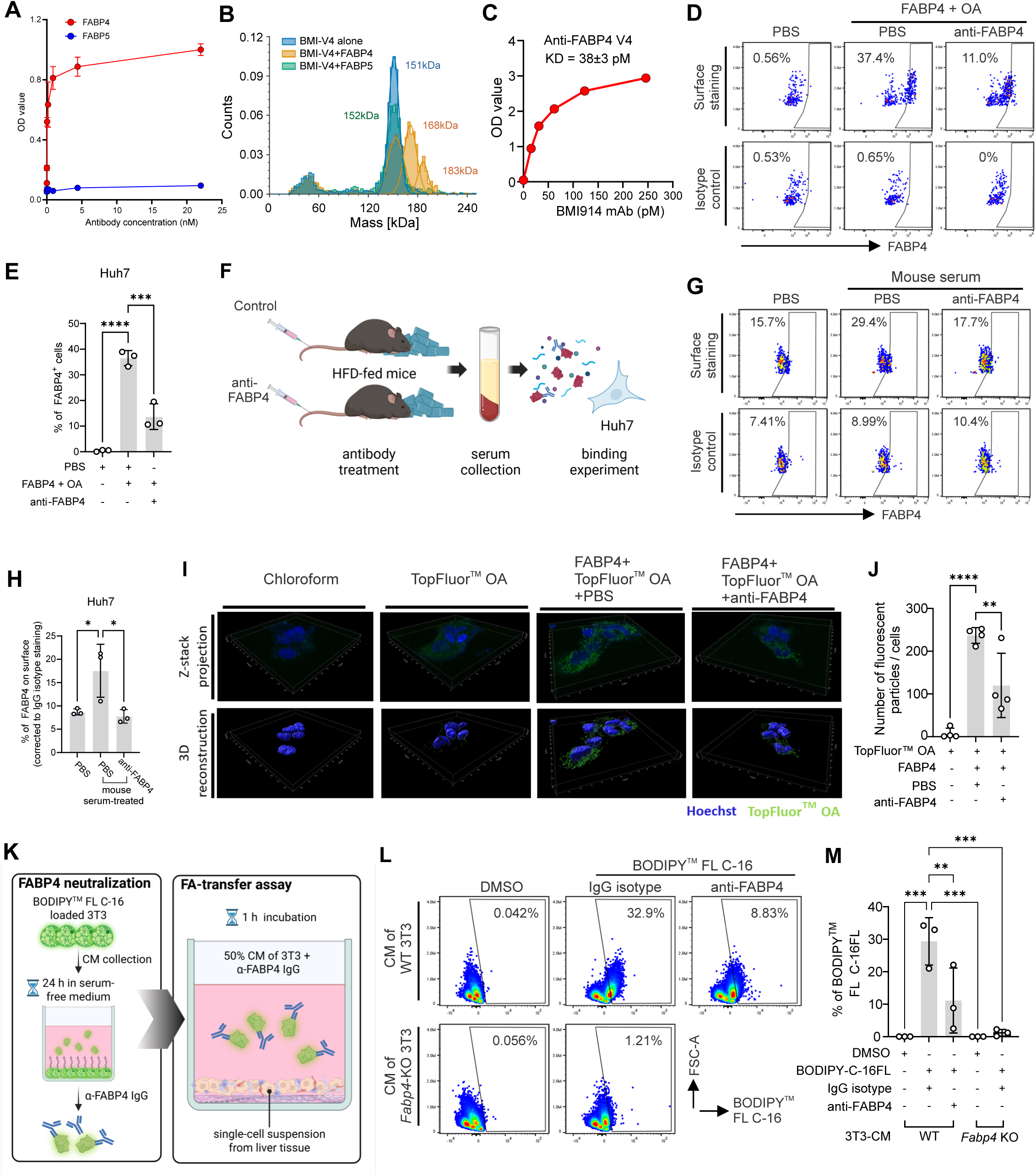
A high-affinity humanized anti-FABP4 antibody blocks FABP4 binding to the cell surface and abrogates fatty acid transfer to hepatocytes. **(A–C)** Biochemical characterization of humanized anti-FABP4 antibody (V4). ELISA binding curves (A) and mass photometry (B) confirmed selective binding of V4 to FABP4 but not FABP5. V4 exhibited picomolar-range binding affinity to FABP4, as determined by equilibrium binding analysis (C). **(D–E)** Representative flow cytometric gating (D) and quantification of surface FABP4 (E) in Huh7 after 15 min of incubation with FABP4-conjugated with OA (1:1; 1 μmol/L) either neutralized with PBS or α-FABP4 IgG (0.5 μmol/L). PBS was used as a vehicle control. IgG isotype control was used as a staining control of surface FABP4. **(F–H)** Schematic *in vitro* FABP4 binding experiment of Huh7 with 0.5% serum that was collected from the HFD-fed mice treated with either PBS or α-FABP4 IgG via tail vein injection (F). Representative flow cytometric gating (G) and quantification of surface FABP4 (H) on Huh7 after 15 min of incubation with 0.5% serum collected from the mice treated with either PBS or α-FABP4 IgG (5 mg/kg; i.v.). PBS was used as a vehicle control. IgG isotype control was used as a staining control of surface FABP4. **(I–J)** Representative Z-projection (Top row) and 3D reconstruction (bottom row) (I) of Huh7 after 15 min incubation with either free form or FABP4-conjugated form (1:1) of TopFluor^TM^ OA (0.5 μmol/L). For FABP4 neutralization, FABP4-conjugated TopFluor^TM^ OA were initially neutralized with either PBS or α-FABP4 IgG (0.25 μmol/L) prior to incubation. Quantification of the number of discontented components of TopFluor^TM^ OA per number of nuclei (Hoechst) from the 3D reconstructed image was calculated (J). Chloroform was used as a vehicle control. **(K)** Schematic of fluorescence FA transfer to primary liver cells using CM collected from the differentiated 3T3 cells. The CM was collected in serum-free RPMI for 16 hours of culture from differentiated 3T3-loaded BODIPY^TM^ FL C-16. The CM is initially neutralized with α-FABP4 IgG prior to incubation with primary liver cells that were prepared by in situ enzymatic digestion. (**L-M**) Representative flow cytometric gating (L) and Quantification of BODIPY^TM^ FL C-16 fluorescence intensity (M) in primary hepatocytes after 1 h of coculture with 50% CM obtained from BODIPY^TM^ FL C-16-loaded differentiated 3T3, as shown in panel K. For FABP4 neutralization, the CM were initially neutralized with either PBS or α-FABP4 IgG (10 nM) prior to incubation. DMSO was used as a vehicle control. Data are presented as mean ± SEM. Statistical testing was performed using one-way ANOVA. *, p < 0.05; **, p < 0.01; ***, p < 0.001; ****, p < 0.0001; ns, nonsignificant.

Because circulating FABP4 binds to hepatocytes to facilitate FA transfer, we first evaluated whether the V4 antibody could block FABP4 surface binding. Exposure of Huh7 cells to soluble FABP4 resulted in detectable surface binding in approximately 40% of cells. Strikingly, V4 antibody treatment significantly reduced FABP4 binding (Figure 6D, 6E). Consistent results were observed in Hepa1-6 cells (Figure S6E, S6F). To verify antibody efficacy in a more physiological context, we administered V4 mAb to HFD-induced obese mice and assessed circulating FABP4 binding using collected serum (Figure 6F). Compared with the control group, V4 mAb markedly reduced circulating FABP4 binding to Huh7 cells (Figure 6G, 6H).

We further examined whether V4 mAb inhibited FABP4-mediated FA uptake. Confocal imaging demonstrated that FABP4-mediated OA uptake by Huh7 cells was significantly suppressed by V4 mAb treatment (Figure 6I, 6J). Flow cytometric analysis confirmed that V4 mAb also inhibited FABP4-mediated OA and PA uptake by Hepa1-6 cells (Figure S6G-S6J). Importantly, using primary mouse hepatocytes isolated via *in situ* enzymatic digestion (Figure 6K), we demonstrated that V4 mAb effectively blocked adipocyte-derived FABP4-mediated FA uptake (Figure 6L, 6M). Collectively, these results indicate that we have generated a high-affinity humanized anti-FABP4 mAb that potently blocks FABP4 binding to hepatocytes and suppresses FABP4-mediated FA uptake in both hepatocyte cell lines and primary hepatocytes, supporting FABP4 as a viable therapeutic target for hepatic steatosis.

### Humanized anti-FABP4 monoclonal antibody attenuates obesity-associated hepatic steatosis in multiple animal models

To evaluate whether the humanized anti-FABP4 V4 mAb mitigates obesity-associated hepatic steatosis *in vivo*, we first assessed its therapeutic efficacy in HFD-induced obese mice. C57BL/6 mice were fed by a HFD (60% fat) for 3 months to induce obesity and subsequently treated weekly with either anti-FABP4 antibody or control IgG (Figure 7A). Body weight remained comparable between treatment groups (Figure S7A), indicating that anti-FABP4 mAb does not alter obesity *per se*.

**Figure 7.**
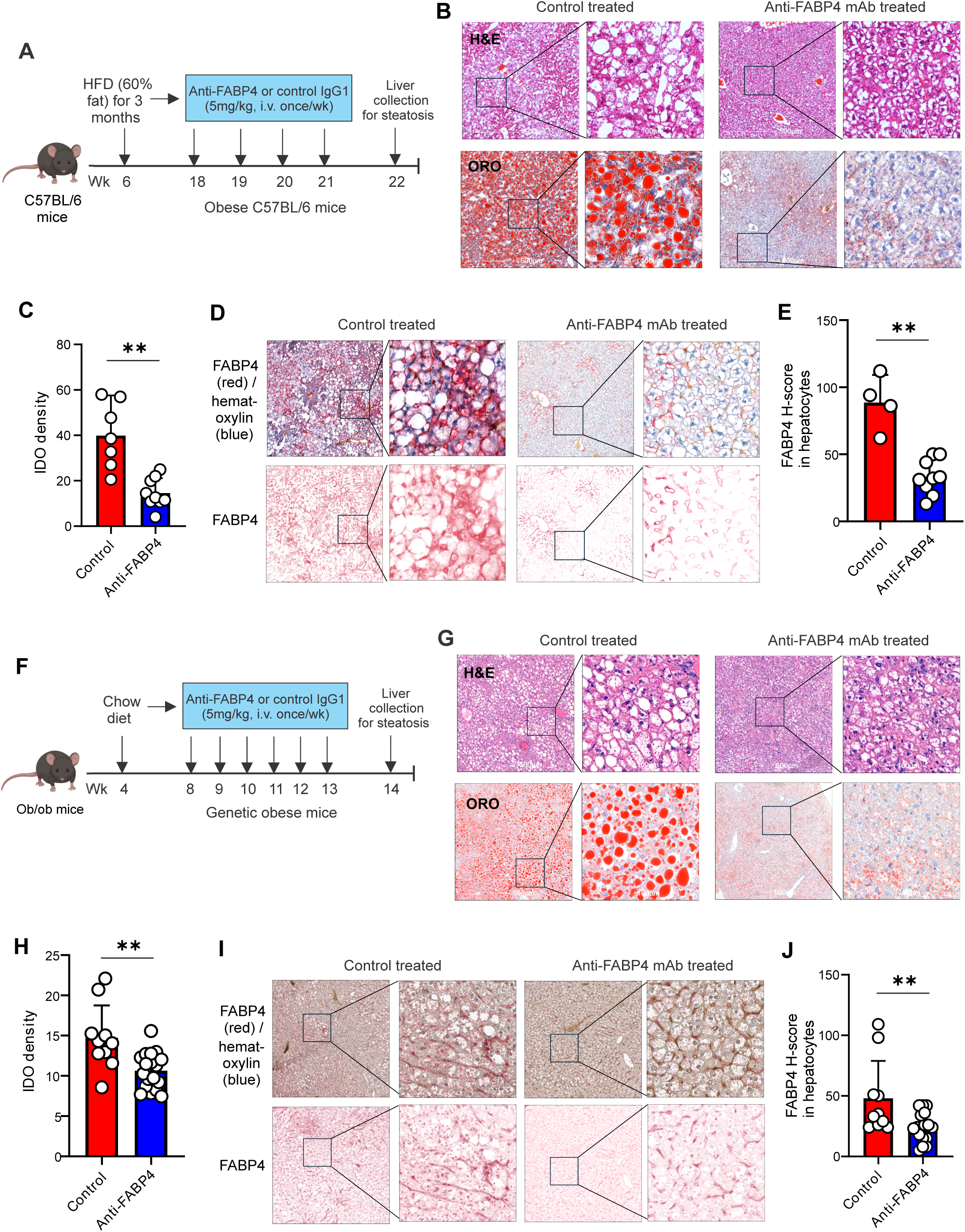
Anti-FABP4 monoclonal antibody treatment reduces hepatic steatosis in diet-induced and genetic obesity models. **(A)** Experimental schematic of the diet-induced obesity model. C57BL/6 mice were fed a high-fat diet (HFD; 60% fat) for 3 months to induce obesity and then treated weekly with anti-FABP4 monoclonal antibody (mAb; 5 mg/kg, i.v.) or control treated Livers were collected for steatosis analyses. **(B)** Representative hematoxylin and eosin (H&E) and Oil Red O (ORO) staining liver sections from control treated and anti-FABP4 mAb–treated HFD-fed mice. **(C)** Quantification of hepatic lipid accumulation based on ORO integrated optical density (IOD) as shown in panel B. **(D)** Representative immunohistochemistry (IHC) staining of FABP4 (red) with hematoxylin counterstain (blue) in liver sections from control treated and anti-FABP4 mAb–treated HFD-fed mice. **(E)** Quantification of hepatocytic FABP4 expression based on H-score analysis in control treated and anti-FABP4 mAb–treated HFD-fed mice. **(F)** Experimental schematic of the genetic obesity model. Leptin-deficient *ob/ob* mice were treated weekly with anti-FABP4 monoclonal antibody (5 mg/kg, i.v.) or control from 8 to 13 weeks of age, followed by liver collection. **(G)** Representative H&E- and ORO-stained liver sections from control treated and anti-FABP4 mAb–treated *ob/ob* mice. **(H)** Quantification of hepatic lipid accumulation based on ORO integrated optical density (IOD) as shown in panel **G**. **(I–J)** Representative IHC staining of FABP4 protein accumulation (red) in hepatocytes of from control treated and anti-FABP4 mAb–treated *ob/ob* mice. **(J)** Quantification of hepatocytic FABP4 H-score is shown in panel **J**. Data are presented as mean ± SEM. Statistical significance was determined by Student’s *t*-test (\*\**p*<0.01).

Despite no effect on body weight, anti-FABP4 mAb treatment markedly reduced macrovesicular hepatic steatosis and neutral lipid accumulation as assessed by histological analysis and ORO staining (Figure 7B). Quantification of ORO staining demonstrated a significant reduction in hepatic lipid deposition in antibody–treated mice compared with controls (Figure 7C). IHC analysis further showed a substantial decrease of FABP4-positive hepatocytes following antibody treatment (Figure 7D,7E). In contrast, FABP5 expression in hepatocytes remained low and unchanged across all groups (Figure S7B, S7C), supporting the specificity of the antibody.

To extend these findings to genetic-induced obesity, we next evaluated the therapeutic efficacy of anti-FABP4 mAb in leptin-deficient *ob/ob* mice, a well-established model of hepatic steatosis^35^. *Ob/ob* mice were treated weekly with anti-FABP4 mAb or control IgG from 8 to 13 weeks of age (Figure 7F). Consistent with the diet-induced obese model, antibody treatment did not affect body weight of *ob/ob* mice (Figure S7D) but significantly reduced hepatic lipid vacuolization and accumulation as assessed by histology and ORO staining (Figure 7G). Quantitative analysis confirmed a robust reduction in hepatic lipid deposition in antibody-treated mice (Figure 7H). IHC analysis further demonstrated decreased FABP4 staining in hepatocytes following anti-FABP4 mAb treatment (Figure 7I, 7J). Consistently, anti-FABP4 mAb treatment did not alter FABP5 expression in hepatocytes (Figure S7E, S7F), underscoring the specificity of FABP4 inhibition.

Collectively, these results demonstrate that therapeutic targeting of circulating FABP4 with a humanized monoclonal antibody selectively blocks FABP4 engagement with hepatocytes and significantly attenuates hepatic steatosis in both diet-induced and genetic models of obesity, supporting FABP4 blockade as a promising therapeutic strategy for obesity-associated hepatic steatosis.

## Discussion

Obesity is a main driver of hepatic steatosis and a critical determinant of progression along the MASLD spectrum^36^. Although stable isotope–tracer studies have established adipose tissue–derived fatty acids as the dominant source of hepatic lipid accumulation in obesity^6, 7^, the molecular mechanisms governing adipose–liver lipid transfer have remained incompletely understood. In this study, we identify circulating FABP4 as a central mediator of obesity-associated hepatic steatosis and demonstrate that selective neutralization of extracellular FABP4 effectively disrupts adipocyte–hepatocyte lipid crosstalk and attenuates disease in multiple *in vivo* models.

Our findings clarify a longstanding discrepancy between transcriptomic and protein-level observations of FABP4 in the liver^21^. In both human and mouse tissues, hepatocytes lack endogenous FABP4 transcription under physiological conditions, including in individuals with obesity. Nevertheless, FABP4 protein accumulates within hepatocytes in steatosis and cirrhosis, strongly indicating an extrinsic origin. These observations provide direct evidence that FABP4 detected in hepatocytes is acquired from the circulation rather than produced locally, thereby redefining the source and context of FABP4 activity in hepatic disease.

Genetic loss-of-function experiments further establish FABP4 as a nonredundant mediator of obesity-induced hepatic lipid accumulation. Whole-body deletion of FABP4 nearly abolished hepatic steatosis in HFD-induced obese mice without altering body weight, adipocyte morphology, or systemic lipid availability. These findings demonstrate that FABP4 promotes hepatocellular lipid accumulation through mechanisms distinct from global changes in adiposity or lipid supply. Importantly, conditional deletion studies revealed that adipocytes are the dominant source of pathogenic FABP4 in this context, whereas FABP4 expression in macrophages or endothelial cells was dispensable for obesity-associated hepatic steatosis. This cell-specific dissection refines prior models that emphasized intracellular FABP4 signaling in hepatic non-parenchymal cells and instead highlights a previously underappreciated endocrine function of adipocyte-derived FABP4.

Mechanistically, we demonstrate that circulating FABP4 directly binds to hepatocytes and facilitates the uptake of both saturated and unsaturated FAs. This process operates independently of hepatocytic FABP4 expression and provides an efficient mechanism for FA delivery during obesity, a state in which FABP4 secretion from hypertrophic adipocytes is increased due to insulin resistance and enhanced lipolytic signaling^18, 19, 37^. Our coculture systems and primary hepatocyte experiments establish that adipocyte-derived FABP4 is both necessary and sufficient to mediate FA transfer to hepatocytes, supporting a model in which FABP4 functions as a soluble lipid carrier bridging adipose tissue and liver.

These mechanistic insights have direct therapeutic implications. Previous strategies to target FABP4 relied primarily on small-molecule inhibitors, including BMS309403, designed to block intracellular FABP4 activity^38, 39^. Although effective in experimental models, these approaches have faced translational limitations, including off-target effects, limited selectivity among FABP family members, and systemic toxicity^24^. Our findings suggest that these challenges may arise, at least in part, from targeting intracellular FABP4 rather than the pathogenic circulating pool that mediates inter-organ lipid flux.

By selectively targeting extracellular FABP4, we introduce a conceptually distinct therapeutic strategy. We developed a high-affinity humanized monoclonal antibody that neutralizes circulating FABP4 without affecting related FABP family members or intracellular FABP4 functions. This antibody effectively blocks FABP4 binding to hepatocytes, suppresses FABP4-mediated fatty acid uptake, and significantly reduces hepatic steatosis in both diet-induced and genetic models of obesity. Notably, these effects occur without altering body weight, highlighting the potential to decouple hepatic benefit from weight loss and suggesting a favorable safety profile.

The efficacy of FABP4 neutralization across multiple obesity models supports circulating FABP4 as a disease-modifying target rather than a secondary biomarker of metabolic dysfunction. Antibody-based inhibition offers additional translational advantages, including high specificity, prolonged half-life, and selective engagement of the extracellular compartment^40, 41^. Given the strong association between circulating FABP4 levels and MASLD severity in humans^13, 42, 43^, FABP4-neutralizing antibodies may enable both therapeutic intervention and patient stratification (e.g. post-transplant hepatic steatosis^44, 45^) in future clinical studies.

In summary, our work identifies circulating FABP4 as a critical molecular link between obesity and hepatic steatosis. We demonstrate that adipocyte-derived FABP4 directly binds hepatocytes to mediate FA transfer, driving hepatic lipid accumulation independently of *de novo* lipogenesis or systemic lipid availability. Targeting this extracellular pathway with a high-affinity humanized monoclonal antibody effectively attenuates hepatic steatosis in preclinical models, establishing circulating FABP4 as a promising therapeutic target for obesity-associated MASLD.

### Limitations

Several limitations of this study should be acknowledged. First, although we demonstrate robust therapeutic efficacy of FABP4 neutralization in multiple mouse models of obesity-associated hepatic steatosis, the translational relevance to human MASLD will require validation in clinical settings, particularly given interspecies differences in lipid metabolism and FABP4 regulation.

Second, while our data establish circulating FABP4 as a key mediator of fatty acid transfer to hepatocytes, the precise hepatocyte surface receptors or binding partners responsible for FABP4 engagement remain to be identified^27, 46, 47^ and may offer additional therapeutic opportunities. Third, this study focused primarily on hepatic steatosis; whether long-term FABP4 blockade can mitigate downstream inflammatory and fibrotic stages of MASLD progression warrants further investigation. Finally, although antibody-based targeting selectively neutralizes extracellular FABP4, potential effects on FABP4 signaling in other metabolic organs and during chronic treatment will need to be carefully evaluated in future safety and pharmacokinetic studies.

## Acknowledgement

We thank the Comparative Pathology Laboratory and Histology Research Laboratory at the University of Iowa for performing liver section and histological staining, and Proteomic facility at the University of Iowa for mass spectrometry analysis. We also thank Dr. Ling Yang from the University of Iowa for the technical support to this study. B.L. thank the funding support from NIH grants R01AI137324, R01CA180986, U01CA272424, the Fraternal Order of Eagles Diabetes Research Center Bridge to the Cure, and the University of Iowa Research Startup Funds.

## Author contributions

X. J., A.A., J.Y., Z.W., S.L., J.S., J. H. performed experiments and analyzed the data. X.H. helped with mouse colony maintenance and high fat diet-induced obese mouse models. H.T. and B.C. did liver tissue gene analysis. Z.X, and N.S. helped with antigen/antibody binding analysis. S.S. provided intellectual inputs and helped with paper-writing. B.L. designed experiments and wrote the paper.

## STAR Methods

### RESOURCE AVAILABILITY

#### Lead contact

Further information and requests for reagents may be directed to and will be fulfilled by the Lead Contact, Bing Li.

#### Material availability

All unique/stable reagents generated in this study are available from the lead contact with a completed materials transfer agreement.

**Key Resources Table** (see attached)

#### Experimental Models and Study Participant Details

##### Cell lines

Hepa 1-6 (RRID: CVCL_0327), Huh7 (CVCL_2957; Sigma), and 3T3 (RRID: CVCL_0594) were purchased from ATCC and Sigma. For Hepa 1-6 and Huh7 cell lines, cells were cultured in DMEM supplemented with 10% FBS and 50 μg/mL gentamycin sulfate solution (IBI Scientific) at 37℃ in an incubator with 5% CO_2_. Whereas 3T3 cells were cultured in RPMI supplemented with 10% FBS and 50 μg/mL gentamycin sulfate solution. *Fabp4*-KO 3T3 and WT 3T3 cell lines were purchased from Ubigene, and knockout cell lines were generated with CRISPR-U technology. The mRNA and protein level of the *Fabp4*-KO 3T3 cell line was validated from the cell lysate. All cell lines were cultured for less than two months after thawing. Mycoplasma testing was not provided by the vendor.

##### Patient samples

Human liver tissue microarrays were obtained from a commercial source (LV1201b; TissueArray.com). The LV1201b microarray contained liver tissue specimens representing multiple disease states, including normal liver, fatty degeneration, cirrhosis, hepatitis, hepatocellular carcinoma, cancer-adjacent liver tissue, cysts, cavernous hemangioma, and echinococcosis. The array includes 118 patient cases across 120 cores, with one core per case. Available clinicopathological annotations include pathological grade, TNM stage, and clinical stage, as provided by the manufacturer. Unstained sections were used for immunohistochemical analyses according to standard protocols.

##### Mouse and diets

Fabp4-/- mice, including whole-body and tissue specific knockout mice, and wild-type littermate controls on a C57BL/6 background were maintained and bred under specific pathogen–free conditions in the animal facility at the University of Iowa, with ad libitum access to water and standard chow. All animal procedures were approved by the University of Iowa Institutional Animal Care and Use Committee (IACUC; protocol #1042385) and were conducted in accordance with NIH and institutional guidelines. All mice (5-6 weeks old) were randomly assigned to experimental groups (7–10 mice per group) and fed either a low-fat diet (LFD; 10 kcal% fat) or a high-fat diet (HFD; 60 kcal% fat) for 3 months prior to tissue collection. Body weight was measured every other week throughout the feeding period.

To generate cell type–specific *Fabp4* knockout mice, *Fabp4* floxed mice (*Fabp4*f/f) were generated by flanking exons 2 and 3 of the *Fabp4* gene with loxP sites *Fabp4*^f/f^ Adipoq-Cre, Fabp4^f/f^ CSF1R-Cre, Fabp4^f/f^ Tek-Cre info: we generated *Fabp4* floxed mice (*Fabp4*^f/f^) by flanking exons 2 and 3 of the *Fabp4* gene with LoxP sites (Figure S3A). *Fabp4*^f/f^ mice were subsequently crossed with Adipoq-Cre, CSF1R-Cre, or Tek-Cre transgenic lines to generate adipocyte-, macrophage-, or endothelial cell-specific Fabp4 knockout mice, respectively. Mice were fed on 60 kcal% HFD or LFD for 3 months before tissue collection. To analyze FABP4 expression in adipocytes, endothelial cells and macrophages, a single-cell suspension was prepared from adipose tissue, liver, and peritoneal cavity from CO_2_-euthanized mice. Briefly, adipose tissue and liver were minced and dissociated into 2 to 4 mm fragments, followed by incubation at 37°C in a digital incubator for 45 minutes on a microplate shaker in RPMI 1640 medium containing 0.5 mg/mL collagenase type II, 0.02 mg/mL DNase I, 0.2 mg/mL hyaluronidase, and 5% FBS. The cell suspension was acquired by treating the fragments with thorough vortexing in a 50 mL tube, followed by filtering through a 40 μm cell strainer into a 50 mL tube. For peritoneal macrophages, 6mL of PBS was injected into the peritoneal cavity left to sit for 3 min. The skin was carefully cut open without damaging the peritoneum, the single cell suspension waw acquired by aspirating the PBS from the peritoneal cavity using a 10mL syringe with a 25G needle, then transferred to a 15mL conical tube. For intracellular FABP4 staining, cells stained by surface markers underwent the intracellular staining process using anti-mouse FABP4 antibody for 1.5 h at 4°C, followed by secondary staining of Donkey-anti-goat IgG-AF647 antibody. Cells were acquired by Cytek Aurora Flow Cytometer.

#### Method Details

##### Antibody treatment

Wild-type mice were fed a high-fat diet (HFD; 60 kcal% fat) for 3 months starting at 1 month of age. Mice were then randomly assigned to experimental groups (control IgG or anti-FABP4 monoclonal antibody) and treated once weekly with antibody (5 mg/kg, i.v.) for 4 weeks. Liver tissues were collected at the end of the treatment period for subsequent analyses.

Leptin-deficient *ob/ob* mice were purchased from The Jackson Laboratory and allowed to acclimate for 4 weeks. Mice were then randomly assigned to experimental groups (control IgG or anti-FABP4 monoclonal antibody) and treated once weekly with antibody (5 mg/kg, i.v.) for 6 weeks, followed by tissue collection.

##### Serum collection

Blood serums were collected from mice via submandibular vein collection. A blood volume of 200 μL was collected from a single mouse. The blood was allowed to clot at room temperature, followed by centrifugation at 10,000g for 5 minutes. The supernatants were collected for the designated assay.

##### Isolation of liver primary cells

High-yield hepatocytes isolation from C57Bl/6j mouse liver was done according to the previously published protocol with several adjustments ^48^. Digestion enzymes were prepared in digestion buffer containing 0.5 mg/mL collagenase type II (Worthington Biochemical), 0.2 mg/mL hyaluronidase (Sigma), and 0.02 mg/mL DNase I (Sigma). After the incision of the portal vein is made, perfusion buffer was infused through the vena cava for 5 min to remove the blood from the liver, followed by a 10 min infusion of digestion buffer. The downstream procedures were done following the published protocol. The obtained single-cell suspension was kept on ice for the designated assay.

##### Fluorescence fatty acid uptake

Fatty acid uptake at the cellular level was measured with BODIPY^TM^ FL C-16 (ThermoFisher) and TopFluor^TM^ OA (Avanti Research) were used as fluorescence-labelled FAs. The BODIPY^TM^ FL C-16 stock was reconstituted at 2 mM in DMSO, whereas TopFluor^TM^ OA was provided in Chloroform at a 1.63 mM concentration. An equal molar concentration of Fluorescence-labelled FA and FABP4 protein (1:1) was incubated in RPMI for 5 min at 37 ℃ to conjugate FABP4 with the Fluorescence-labelled FA. To measure the FA uptake in the cell lines, a number of 1 x 10^4^ cells/well of Hepa 1-6 or Huh7 were seeded in a 48-well plate and were cultured overnight. Prior to the uptake, the cells were starved in serum-free medium for 3 hours before the uptake began. Both the Free form and FABP4-conjugated form of fluorescence-labelled FA were directly added to the cell culture medium for 15 min at 37 ℃. The cells were then harvested with 0.025% Trypsin for Zombie Violet staining to distinguish between live and dead cell populations. The data acquisition was performed with a Cytek Aurora spectral flow cytometer.

##### Generation of humanized anti-FABP4 mAbs

Anti-FABP4 antibodies was generated and screened as previous described^30^. For humanized antibody production, parental VH and VL sequences were run through a CDR grafting algorithm to transfer the CDRs from the original framework onto the most matched human germline sequences. To ensure that no highly undesirable sequence liabilities were introduced into the humanized sequences, identified high-risk motifs were removed through mutagenesis. A total of 16 antibody variants composed of different pairings of 4 humanized heavy chains and 4 humanized light chains were generated using CHO mammalian cells. All humanized antibodies were purified from CHO cells with high purify and low endotoxin (<0.05EU/mg) for *in vivo* studies.

##### Surface FABP4 binding assay

Stearic acid (SA) and Oleic acid (OA) were purchased from Nu-Chek Prep, Inc., and diluted in EtOH at 50 mM as a stock concentration. The free fatty acid (1μmol/L) was then conjugated with FABP4 protein at equal molar concentration (1:1) at 37 ℃ for 5 min. To measure the FABP4 binding on the cell surface, 1x10^4^ cells of Hepa 1-6 or Huh7 were seeded in a 48-well plate and cultured overnight. The cells were then serum-starved for 3 hours prior to the binding assay. FABP4-conjugated FA was added to the culture medium for 15 min at 37 ℃. For quantifying the percentage of mouse serum-derived FABP4 that binds to the cell surface, 0.5% of serum was added to the serum-free culture medium and cultured for 15 min at 37 ℃. The cells were then harvested with 0.025% Trypsin and stained on ice for 30 min with Goat α FABP4 and Zombie Violet, followed by secondary antibody staining. For the isotype control staining, Normal Goat IgG (R&D Systems) was used. The whole procedure of surface staining was maintained on ice. The data acquisition was performed with a Cytek Aurora spectral flow cytometer.

##### In vitro Ab neutralization

To block FABP4 protein functionality, FABP4-conjugated with either FA or fluorescence labelled FA was mixed with α-FABP4 IgG at half of the molar concentration of the FABP4 protein to anti-FABP4 (1:2) for 3 hours at 37℃ before incubating it in the cell culture medium. For neutralizing FABP4 in the CM collected from BODIPY^TM^ FL C-16-loaded differentiated 3T3 cells, 10 nmol/L of α-FABP4 IgG was added to the CM and incubated for 3 hours at 37 ℃. To challenge the antibody efficacy to block FABP4-mediated FA transfer, the incubated mixture of CM-α-FABP4 IgG was incubated with the primary liver cells that were seeded at a density of 10 x 10^4^ cells/well in a 48-well plate with 0.01% collagen pre-coated three hours prior to seeding. After one hour of culture with the CM-α-FABP4 IgG mixture, the primary liver cells were harvested by gentle pipetting and stained for zombie viability marker, followed by fluorescence intensity detection by flow cytometer.

##### Adipogenic 3T3 differentiation and fluorescence-FA loading

3T3 cells were differentiated according to the previously published protocol ^49^. Briefly, for transwell coculture, 3 x 10^4^ of WT and *Fabp4*-KO 3T3 cells were seeded in a 24-well plate and allowed to reach 100% confluency in two days and further cultured in 100% confluency for another two days. During the day of differentiation, differentiation medium one (DM I) containing 0.5 mM IBMX, 1 μM Dexamethasone. 10 μg/mL insulin and 2 μM rosiglitazone were prepared in RPMI supplemented with 10% FBS were added to the 3T3 cells. After two days, the culture medium was replaced with DM II containing 10 μg/mL insulin was prepared in RPMI supplemented with 10% FBS. The DM II was replaced once a day. Afterwards, two days post-differentiation in DM II, a nascent differentiated 3T3 was ready to be loaded with fluorescent FA. To load the nascent differentiated 3T3, the differentiated 3T3 cells were washed twice with 1X PBS to remove excess DM II from the previous culture and cultured in serum-free RPMI with 1 μM of BODIPY^TM^ FL C-16 for 24 hours. After loading 3T3 cells for 24 hours with fluorescence-FA, the cells were then washed twice with 1X PBS and ready for the subsequent transwell coculture. To collect the conditioned medium from 3T3-WT or *Fabp4*-KO, the newly prepared 3T3-loaded fluorescence-FA were immediately cultured in serum-free RPMI for another 24 hours. The supernatant was then collected as conditioned medium and was used for no longer than 3 days for the subsequent indicated assay.

##### Transwell insert coculture

In vitro transwell coculture was performed between fluorescence-FA-loaded 3T3 cells and two models, cell lines and primary hepatocytes. Initially, 3T3 cells were differentiated and loaded with fluorescence FA in a 24-well plate in the lower chamber. Afterwards, the cell medium was washed twice with 1X PBS to remove excess fluorescence-FA in the well and cultured in serum-free RPMI. Concurrently, at the time during the fluorescence-FA loading (one day prior to coculture), Hepa 1-6 or Huh7 were seeded into a 24-well plate insert (pore size: 0.4 μm) with a density of 5 X 10^4^ cells per insert and cultured overnight. On the day of the coculture, the culture medium was removed and washed with 1X PBS twice to remove excess serum in the insert, then replaced with serum-free RPMI before placing the insert with cell lines in the well that contained fluorescence-FA-loaded 3T3. The 3T3-Hepa 1-6 or 3T3-Huh7 coculture was allowed to share the culture medium through the pore for 24 hours. The cells were then harvested with 0.025% trypsin and followed by Zombie viability staining. For 3T3-primary hepatocytes coculture, the newly isolated primary liver cells were seeded in a 24-well plate insert (pore size: 0.4 μm) with a density of 5 x 10^4^ cells per insert, which was pre-coated with 0.01% collagen for three hours prior to seeding. The 3T3-primary hepatocytes were cocultured for 24 hours and harvested for zombie viability staining by gentle pipetting. The intensity of BODIPY^TM^ FL C-16 was acquired using a flow cytometer.

##### Confocal microscopy and 3D reconstruction

Huh7 were seeded with a density of 4 x 10^4^ cells/dish in the 40.4 mm glass-bottom dish and cultured overnight. The cells were serum-starved for 3 hours prior to the TopFluor^TM^ OA uptake assay. Neutralization of FABP4-TopFluor^TM^ OA was prepared as mentioned above and cultured with the cells in serum-free medium for 15 min at 37℃. Afterwards, 5 μg/mL Hoechst was added to the culture for an additional 5 minutes of culture to stain the nuclei. The cells were washed twice with pre-warmed 1X PBS. The imaging was done immediately for no more than 10 min post-washing step with Zeiss LSM 880 confocal microscope. The acquired Z-stack images were subjected to 3D reconstruction with Imaris software. The number of disconnected components of the fluorescence from TopFluor^TM^ OA was calculated and normalized with the number of nuclei. The results were plotted with GraphPad Prism.

##### Histology analysis

For H&E staining, mouse livers were fixed in 4% paraformaldehyde, embedded in O.C.T., and sectioned at 5 µm thickness and were stained with hematoxylin and eosin. Slides were mounted using VectaMount Express Mounting Medium (Vector Laboratories, H-5700-60), and scanned using a Leica Aperio GT 450 slide scanner for subsequent analysis.

For Immunohistochemistry and H-score calculation, multiple-disease human liver tissue microarrays (TMAs; LV1201b) were purchased from Tissue Microarray.com. Paraffin-embedded TMA slides were deparaffinized in xylene and rehydrated through graded ethanol solutions. Antigen retrieval was performed by incubating slides in 10 mM citrate buffer (pH 6.0) at 95 °C for 15 min. Endogenous peroxidase activity was quenched using BLOXALL Endogenous Blocking Solution (Vector Laboratories, SP-6000) for 10 min at room temperature. Slides were incubated with primary antibodies overnight at 4 °C, followed by incubation with appropriate secondary antibodies for 30 min at room temperature. All primary and secondary antibodies used in this study are listed in the Key Resources Table. Signal detection was performed using Vector Red alkaline phosphatase substrate (Vector Laboratories, SK-5100). Slides were counterstained with hematoxylin (Leica Gill III, 3801541) for 10 s at room temperature, mounted using VectaMount Express Mounting Medium (Vector Laboratories, H-5700-60), and scanned using a Leica Aperio GT 450 slide scanner. Whole-slide images of chromogenic sections were generated for subsequent analysis. Slide scans can be made available upon request.

Scanned tissue sections were analyzed using HALO software v3.6.4134.263 (Indica Labs). The percentage and intensity of FABP4- or FABP5-positive cells across whole-slide tissue areas were quantified for H-score calculation^50^. H-scores were calculated using the formula: (percentage of weak intensity × 1) + (percentage of moderate intensity × 2) + (percentage of strong intensity × 3).

For mouse liver immunohistochemistry, tissues were fixed in 4% paraformaldehyde, embedded in O.C.T. compound, and sectioned at 7 µm thickness. Frozen sections underwent endogenous peroxidase quenching and then followed by the same immunohistochemical staining protocol used for human TMA slides. H-score analysis for hepatocytic FABP4 and FABP5 expression in mouse liver sections was performed using the same criteria and calculation method as described for human TMA samples. Single-channel representations of FABP4 and F4/80 staining in the figures were generated by color deconvolution using HALO software (Indica Labs).

##### Oil Red O quantification and IOD density analysis

Oil Red O powder (Alfa Aesar) was solubilized in 100% isopropanol (75 mg in 25 mL) and mixed on an orbital shaker for 30 min to generate a stock solution. For the working solution, the Oil Red O stock was diluted at a ratio of 3 parts Oil Red O to 2 parts ddH₂O and mixed on an orbital shaker for 10 min. The working solution was filtered twice through a 0.45 μm syringe filter and applied to 4% PFA-fixed frozen liver sections. Slides were mounted using VectaMount AQ Aqueous Mounting Medium (Vector Laboratories, H-5501-60) and scanned using a Leica Aperio GT 450 slide scanner. Whole-section imaging of chromogenic slides was performed using the same slide scanner for subsequent quantitative analysis.

Whole-slide images were analyzed using HALO software (v3.6.4134.263; Indica Labs). Lipid-positive regions were identified using a consistent classifier and threshold applied across all samples. The average optical density (OD) of ORO-positive cytoplasmic staining was measured from a representative region of interest (0.8 × 1.1 mm²) to reflect staining intensity. In parallel, the percentage of ORO-positive area was quantified across the entire tissue section. To account for variability in tissue size between sections, an IOD density metric was calculated as the product of the average positive cytoplasmic OD and the percentage of positive area for each slide. This approach integrates both staining intensity and extent of lipid deposition and provides a normalized measure of hepatic lipid accumulation independent of total tissue area.

##### Mouse serum biomarker detection by ELISA

Serum FABP4 levels were measured using a mouse FABP4 ELISA kit. Serum albumin concentrations were determined using a Mouse Albumin ELISA Kit. Serum triglyceride (TAG) levels were measured using the LabAssay™ Triglyceride kit based on an enzymatic colorimetric (GPO-DAOS) method. Serum free fatty acid (FFA) levels were quantified using the Free Fatty Acid Fluorometric Assay Kit. Absorbance was detected using a microplate reader, and analyte concentrations were calculated based on standard curves generated in parallel with each assay. All values were normalized to serum volume and are presented as concentrations.

##### Quantification and statistical analysis

All data were acquired from biological replicates. The statistical analyses were carried out using GraphPad Prism 10. The quantitative data are reported as mean ± SD unless otherwise stated. A two-tailed unpaired Student’s *t*-test was used for comparing a two-group study. For multiple groups comparison, an ordinary one-way ANOVA multiple comparisons with correction by Tukey’s methods were performed. Statistically significant *P* values are reported as ∗ *p* < 0.05, ∗∗ *p* < 0.01 and ∗∗∗ *p* < 0.001. The sample size (n) was reported in each figure legend.

