## Supplementary material for "Targeting Circulating FABP4 Ameliorates Obesity-Associated Hepatic Steatosis": Figure S1-S7

**Supplementary Figure S1-S7**

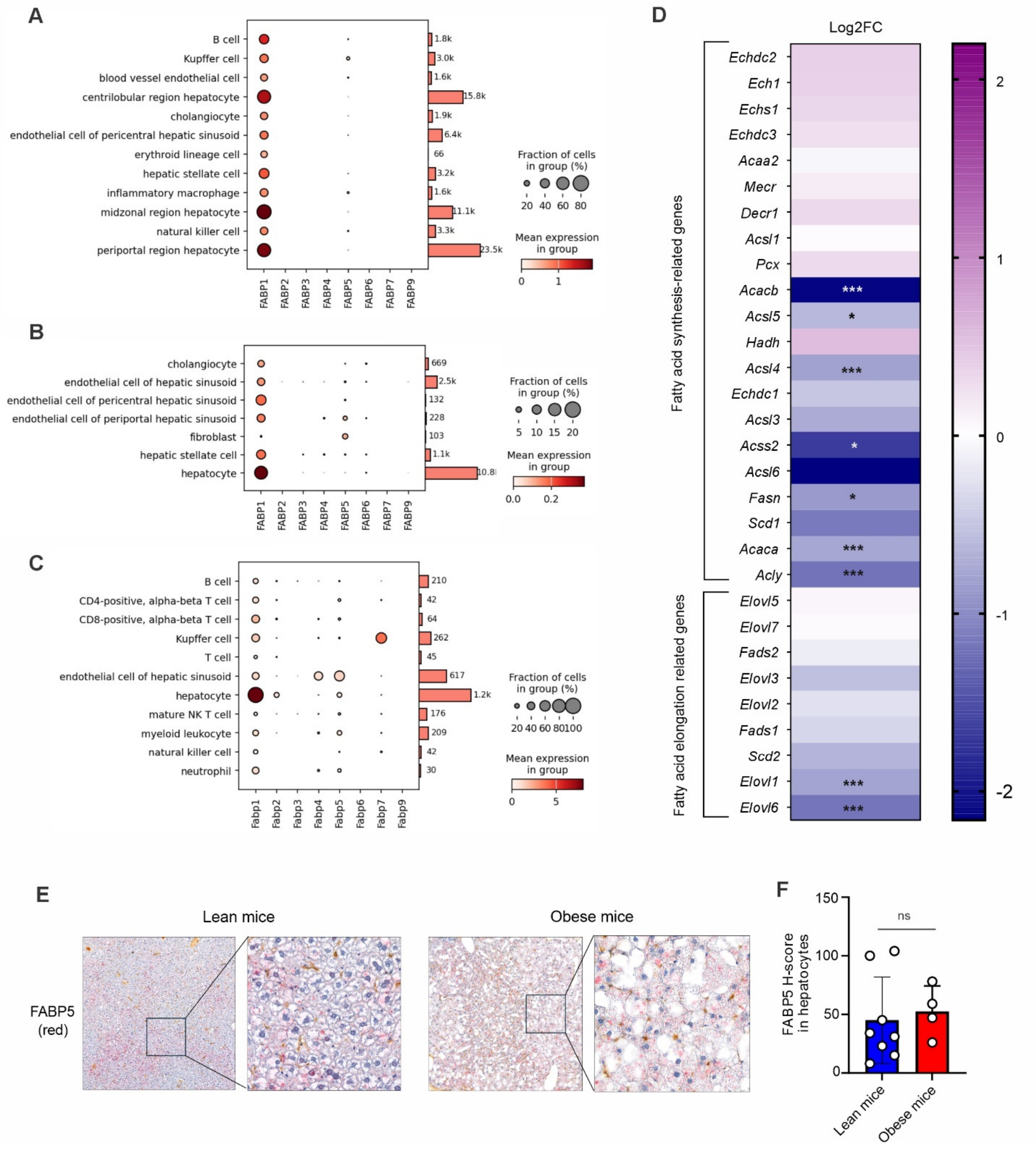


**Figure S1. FABP4 accumulation in hepatocytes occurs independently of transcriptional upregulation**

**(A–C)** Analysis of expression profiles of *FABP* family members across single-cell populations in human (A, B) and mouse liver datasets (C) using BxGenomics.

**(D)** Heatmap showing fold changes in hepatic mRNA expression of fatty acid de novo synthesis- and elongation-related genes in obese mice relative to lean controls.

**(E-F)** Representative IHC images of FABP5 staining (red) in mouse livers of lean and obese mice (E). Quantification of FABP5 H-score in hepatocytes is shown in panel F.

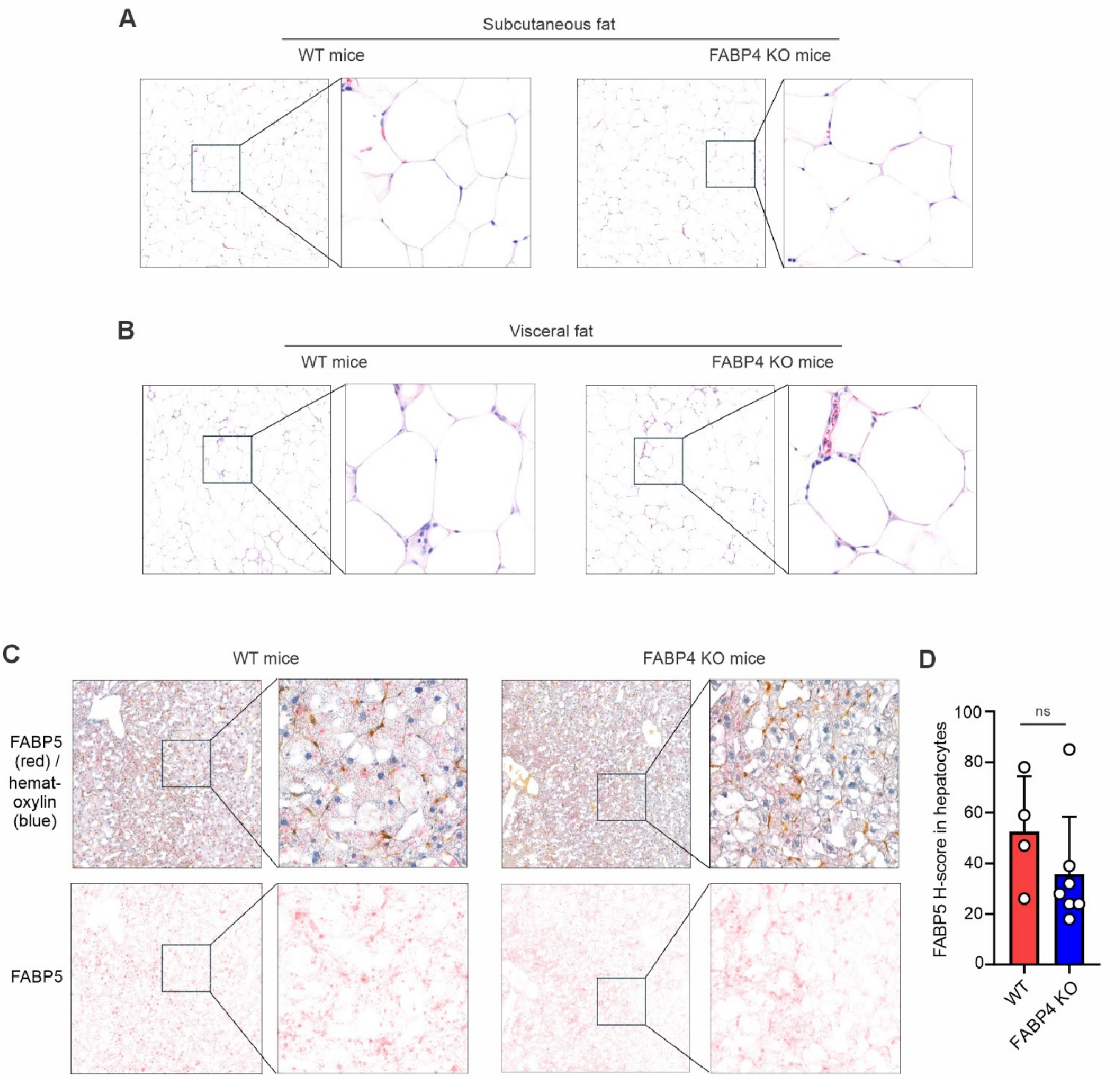


**Figure S2.** ***Fabp4*** **deficiency does not alter adipocyte morphology or systemic lipid availability**

**(A–B)** Representative H&E images of subcutaneous (**A**) and visceral (**B)** adipose tissue from obese WT and *Fabp4* KO mice following HFD feeding for 3-4 months.

**(C–D)** Representative IHC images of FABP5 protein staining (red) and quantification of hepaticFABP5 levels in obese WT and FABP4 KO livers (ns, non-significant).

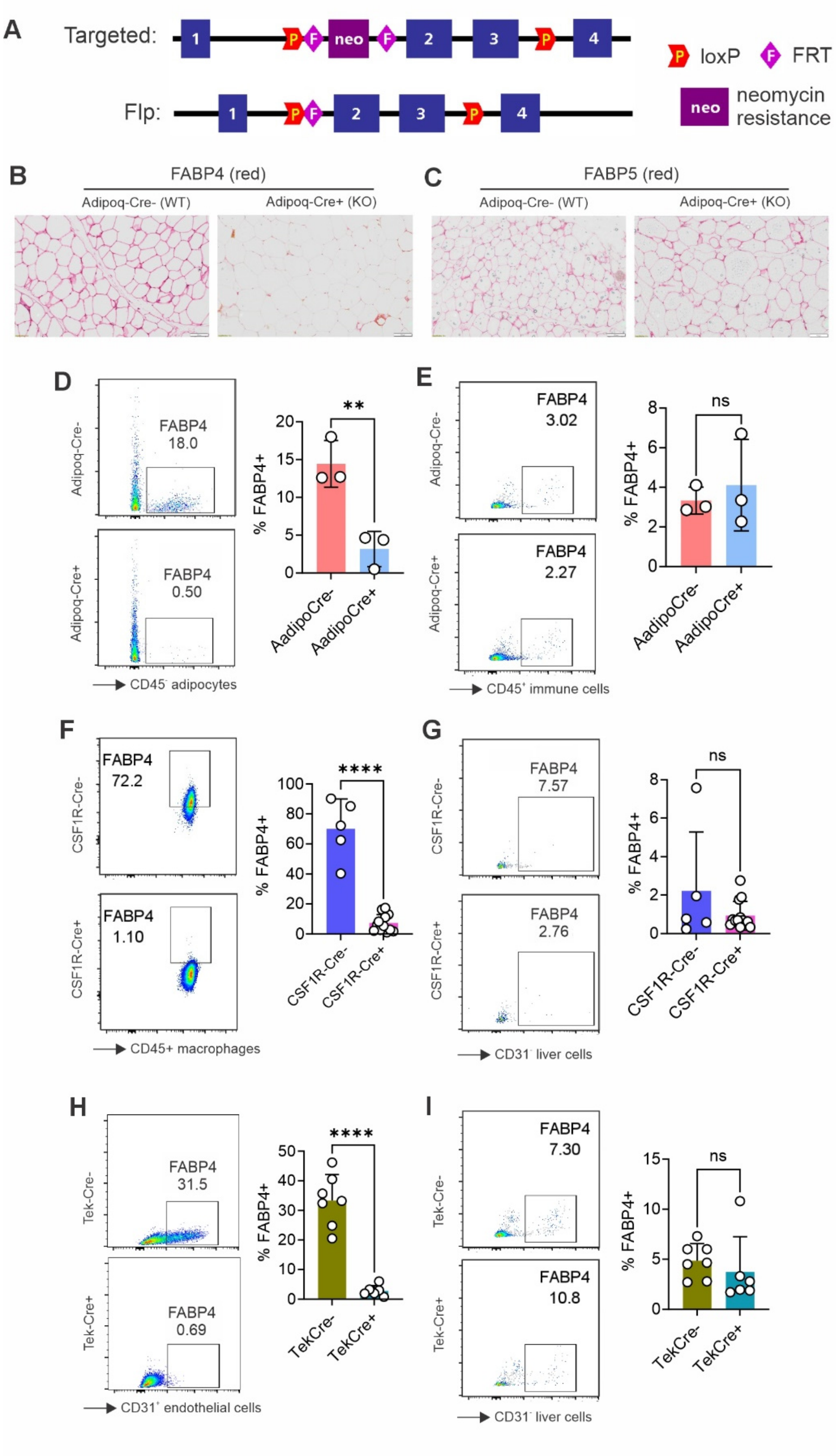


**Figure S3. Generation of tissue specific *Fabp4* knockout mice**

(**A)** strategy of generating *Fabp4*-floxed mice.

(**B**-**C**) Immunohistochemical (IHC) staining of FABP4 (**B**) and FABP5 (**C**) in adipose tissue of *Fabp4*$^{f/f}$ Adipoq-Cre mice.

(**D-E**) Intracellular flow cytometric staining of FABP4 expression and average percentage of FABP4$^{+}$ adipocytes (CD45$^{-}$) (**D**) and CD45$^{+}$ immune cells (F4/80$^{+}$) (**E**) in adipose tissue of *Fabp4*$^{f/f}$ Adipoq-Cre mice.

(**F-G**) Intracellular flow cytometric staining of FABP4 expression and average percentage of FABP4$^{+}$ macrophages (**F**) and CD31$^{-}$ cells (**G**) in livers of *Fabp4*$^{f/f}$ CSF1R-Cre mice.

(**F-G**) Intracellular flow cytometric staining of FABP4 expression and average percentage of FABP4$^{+}$ endothelial cells (CD31$^{+}$) (**F**) and CD31$^{-}$ cells (**G)** in livers of *Fabp4*$^{f/f}$ Tek-Cre mice.

Data are expressed as mean ± SEM; statistical analyses were performed using t-test. **, $p < 0.01$; ****, $p < 0.0001$; ns, nonsignificant.

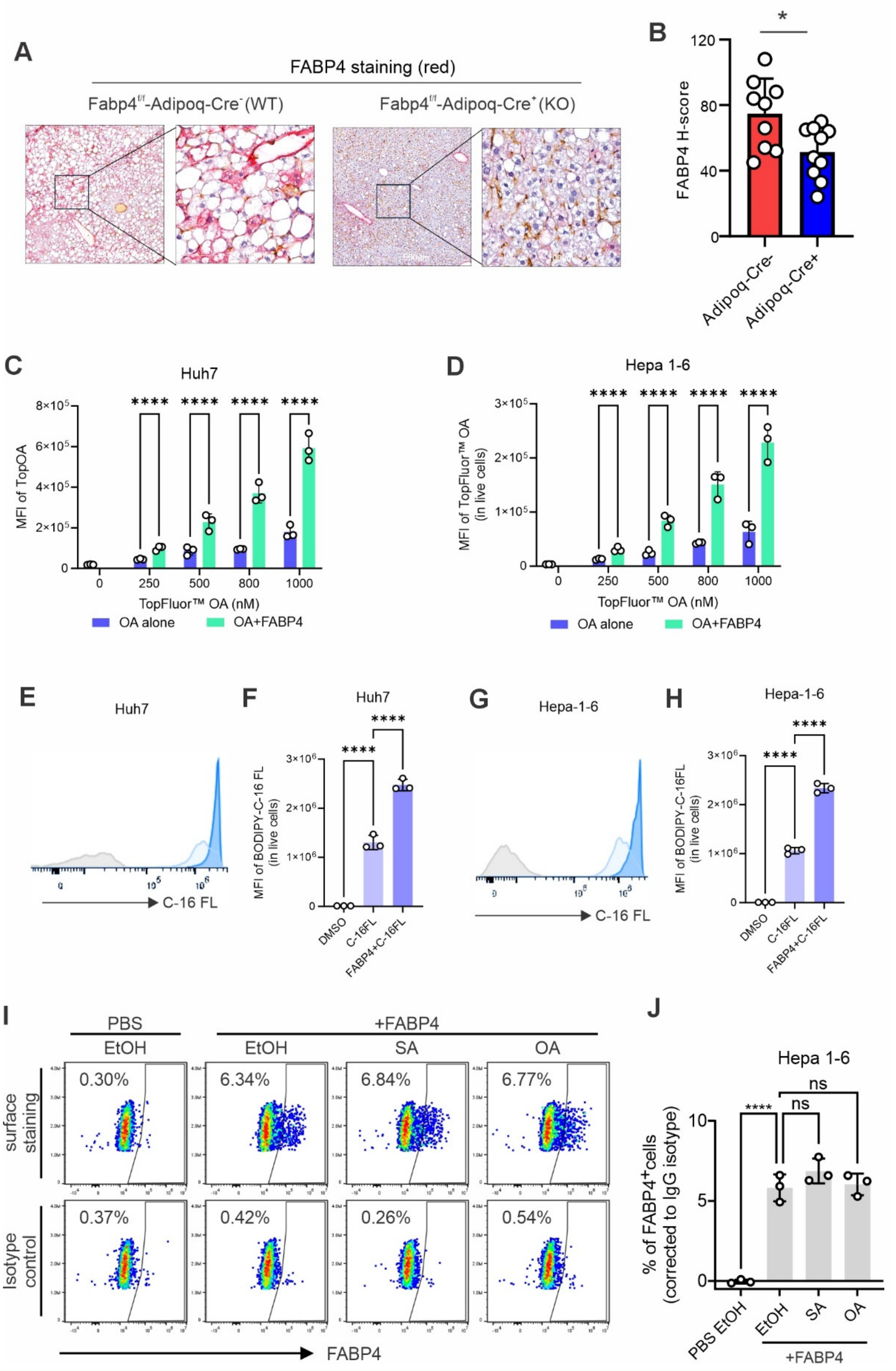

A
FABP4 staining (red)
Fabp4f/f-Adipoq-Cre- (WT)
Fabp4f/f-Adipoq-Cre+ (KO)
B
FABP4 H-score
Adipoq-Cre-
Adipoq-Cre+
C
Huh7
MFI of TopOA
TopFluor™ OA (nM)
OA alone
OA+FABP4
D
Hepa 1-6
MFI of TopFluor™ OA (in live cells)
TopFluor™ OA (nM)
OA alone
OA+FABP4
E
Huh7
C-16 FL
F
Huh7
MFI of BODIPY-C-16 FL (in live cells)
DMSO
C-16FL
FABP4+C-16FL
G
Hepa-1-6
C-16 FL
H
Hepa-1-6
MFI of BODIPY-C-16FL (in live cells)
DMSO
C-16FL
FABP4+C-16FL
I
PBS
+FABP4
EtOH
EtOH
SA
OA
surface staining
Isotype control
0.30%
6.34%
6.84%
6.77%
0.37%
0.42%
0.26%
0.54%
FABP4
J
Hepa 1-6
% of FABP4+cells (corrected to IgG isotype)
PBS EtOH
EtOH
SA
OA
+FABP4
ns
ns

**Figure S4. FABP4 enhances dose-dependent fatty acid uptake and directly binds hepatocytes**

**(A–B)** Representative IHC staining of FABP4 protein accumulation (red) in hepatocytes from *Fabp4*$^{f/f}$-Adipoq-Cre$^{-}$ (WT) and *Fabp4*$^{f/f}$-Adipoq-Cre$^{+}$ (KO) obese mice (A). Quantification of hepatocytic FABP4 H-score is shown in panel B.

**(C–D)** Quantification of TopFluor$^{TM}$ OA fluorescence intensity using flow cytometry in Huh7 (C) and Hepa 1-6 (D) after 15 min incubation of either free form or FABP4-bound form (1:1) TopFluor$^{TM}$ OA at a different concentration (nM) as indicated.

**(E–H)** Representative flow cytometric histogram (E, G) and Quantification of BODIPY$^{TM}$ FL C-16 fluorescence intensity (F, H) in Huh7 (E - F) and Hepa 1-6 (G – H) after 15 min incubation with either free form or FABP4-bound form (1:1) of BODIPY$^{TM}$ FL C-16 (1 µmol/L). DMSO was served as a vehicle control.

**(I-J)** Representative flow cytometric gating (I) and quantification (J) of FABP4 on the surface of Hepa 1-6 after 15 min incubation of FABP4-bound form (1:1) with either EtOH, SA, or OA at 1 µmol/L concentration. PBS and EtOH were used as vehicle control.

Data are expressed as mean ± SEM; statistical analyses were performed using one-way ANOVA (D, F, H) or two-way ANOVA (A, B). ***, $p < 0.001$; ****, $p < 0.0001$; ns, nonsignificant.

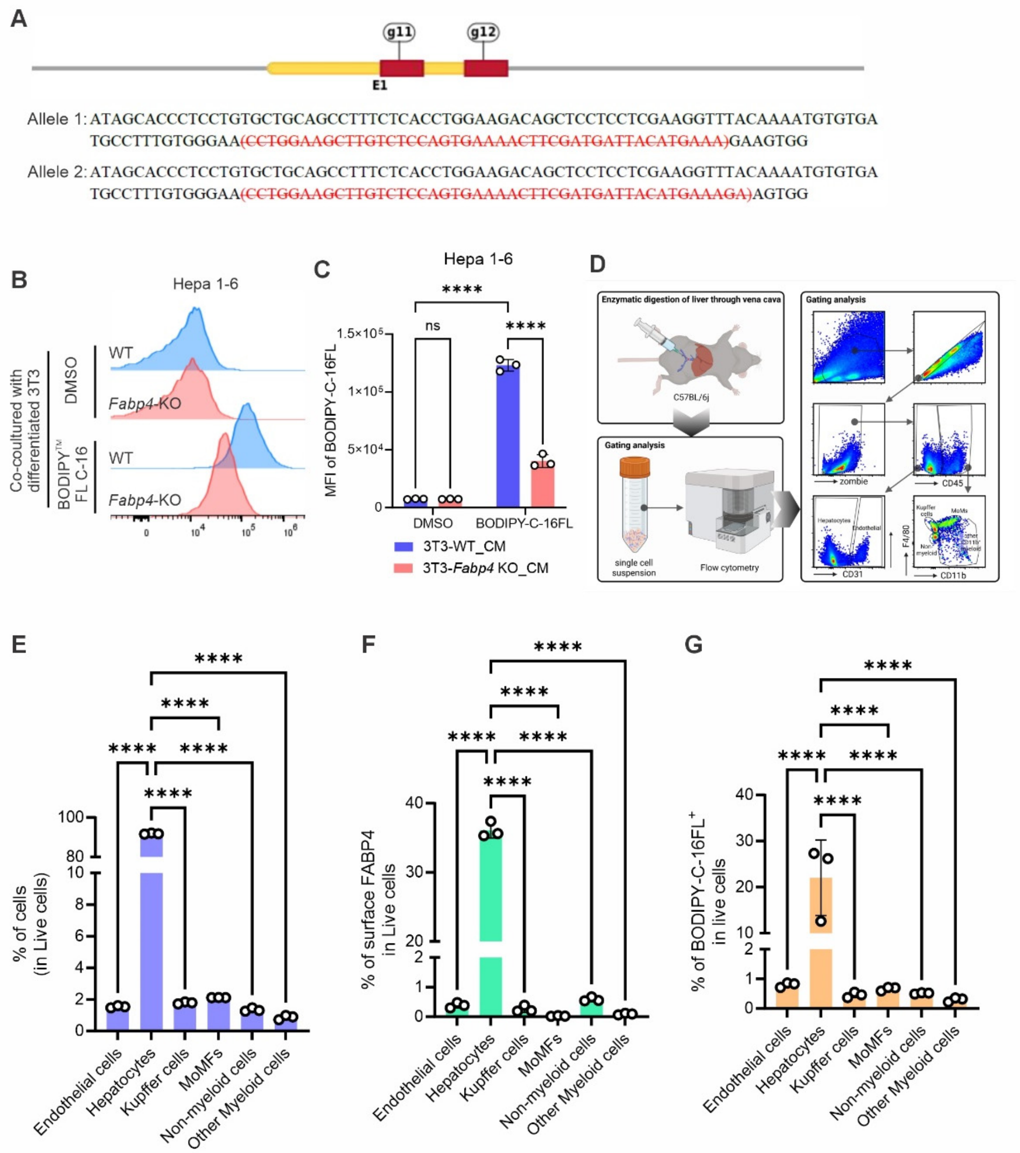


**Figure S5. Generation of *Fabp4*-KO 3T3 cells and FA transfer mediated by FABP4 in primary hepatocytes.**

**(A)** Generation of *Fabp4*-KO in 3T3 cells by CRISPR/Cas9, showing the gRNA flanking the targeted region in exon 1 of *Fabp4* gene. The deletion in both alleles is marked by red strikethrough.

**(B–C)** Representative flow cytometric histogram (B) and Quantification of BODIPY™ FL C-16 fluorescence intensity (C) in Hepa 1-6 after 1 h of coculture with the CM collected from either BODIPY™ FL-C16-loaded WT or *Fabp4*-KO differentiated 3T3. DMSO was used as a vehicle control in both WT and *Fabp4*-KO differentiated 3T3 during the loading

**(D)** Schematic procedure for isolating single cell suspension from liver tissue and gating strategy for flow cytometric analysis.

**(E)** Flow cytometric analysis of the percentage of cell populations from the single cell suspension isolated via *in situ* enzymatic digestion of the primary mouse liver.

**(F-G)** Quantification of surface FABP4 (F) and fluorescence intensity of BODIPY™ FL C-16 (G) in the live cells population of single cell suspension from the liver after 1 h incubation with 50% of conditioned medium (CM) obtained from overnight culture of BODIPY™ FL C-16 (1 μmol/L)-loaded differentiated 3T3 cells in RPMI.

Data are expressed as mean ± SEM; statistical significance was determined using one-way ANOVA. *, $p < 0.05$; ***, $p < 0.001$; ****, $p < 0.0001$; ns, nonsignificant.

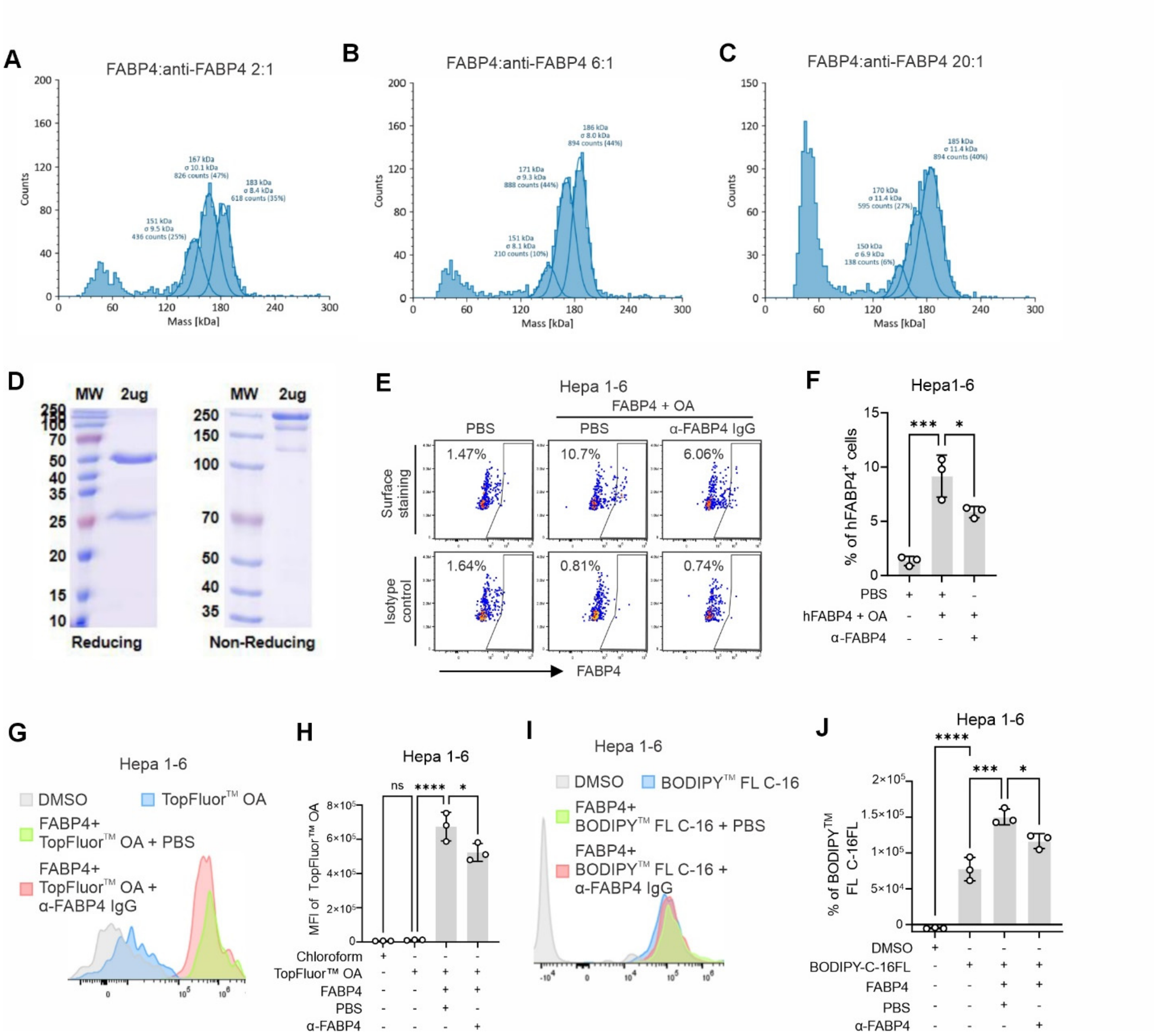

**Figure S6. Biophysical validation and functional confirmation of anti-FABP4–mediated inhibition of fatty acid transfer.**

**(A–C)** Mass photometry of FABP4–antibody complexes at increasing FABP4:V4 molar ratios showing stable complex formation and expected stoichiometry.

**(D)** SDS-PAGE under reducing and non-reducing conditions, confirming the purity and structural integrity of the V4 antibody.

**(E–F)** Representative flow cytometric gating (E) and quantification of surface FABP4 (F) in Hepa 1-6 after 15 min of incubation with FABP4-conjugated with OA (1:1; 1 μmol/L) either neutralized with PBS or α-FABP4 IgG (0.5 μmol/L). PBS was used as a vehicle control. IgG isotype control was used as a staining control of surface FABP4.

**(G–H)** Representative flow cytometric histogram (G) and Quantification of TopFluor$^{TM}$ OA fluorescence intensity (H) in Hepa 1-6 after 15 min of incubation with either free form or FABP4-conjugated form of TopFluor$^{TM}$ OA (1:1; 1 μmol/L), either neutralized with PBS or α-FABP4 IgG (0.5 μmol/L). PBS was used as a vehicle control.

**(I–J)** Representative flow cytometric histogram (I) and Quantification of BODIPY$^{TM}$ FL C-16 fluorescence intensity (J) in Hepa 1-6 after 15 min of incubation with either free form or FABP4-conjugated BODIPY$^{TM}$ FL C-16 (1:1; 1 μmol/L), either neutralized with PBS or α-FABP4 IgG (0.5 μmol/L). PBS was used as a vehicle control.

Data shown as mean ± SEM; statistical tests performed using one-way ANOVA. *, $p < 0.05$; ***, $p < 0.001$; ****, $p < 0.0001$; ns, nonsignificant.

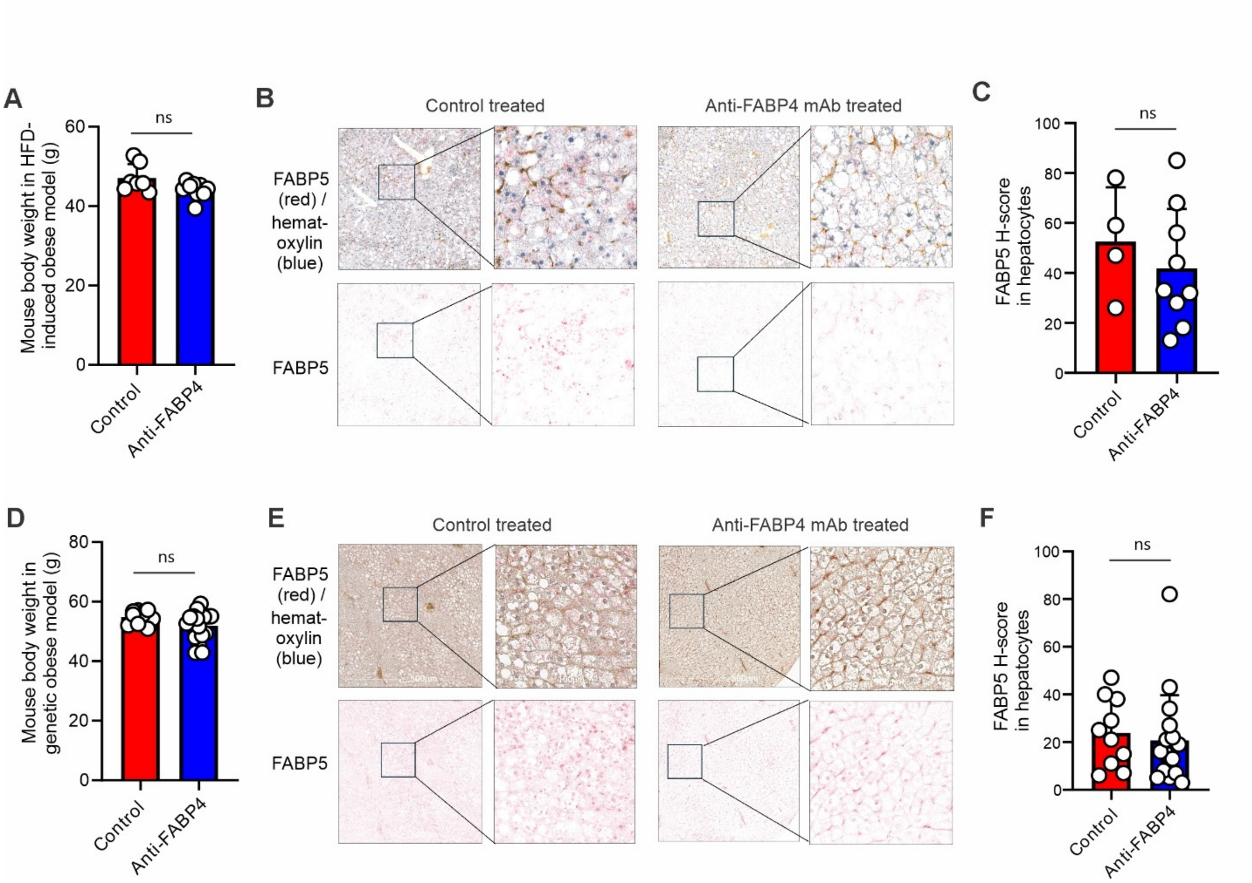


**Figure S7. Anti-FABP4 monoclonal antibody treatment does not alter hepatocytic FABP5 expression**

**(A)** Body weight of high-fat diet (HFD)–induced obese C57BL/6 mice treated with anti-FABP4 monoclonal antibody (mAb) and control treated.

**(B)** Representative immunohistochemistry (IHC) staining of FABP5 (red) with hematoxylin counterstain (blue) in liver sections from control treated and anti-FABP4 mAb–treated HFD-fed mice.

**(C)** Quantification of hepatocytic FABP5 expression based on H-score analysis in control treated and anti-FABP4 mAb–treated HFD-fed mice.

**(D)** Body weight of leptin-deficient *ob/ob* mice treated with control treated or anti-FABP4 monoclonal antibody treated.

**(E)** Representative IHC staining of FABP5 (red) with hematoxylin counterstain (blue) in liver sections from control treated and anti-FABP4 mAb–treated *ob/ob* mice.

**(F)** Quantification of hepatocytic FABP5 expression based on H-score analysis in control treated and anti-FABP4 mAb–treated *ob/ob* treated mice. (please add ‘ns’)

Data are presented as mean ± SEM. Statistical significance was determined by Student’s *t*-test; ns, nonsignificant.
